# Elevated Expression of *MALAT1* Contributes to the Survival of Drug-Tolerant Persister Cells Following Targeted Therapy in Lung Adenocarcinoma

**DOI:** 10.64898/2026.05.07.723110

**Authors:** WJH Davis, M Thompson, SM Farry, C McKinney, G Gimenez, ME Hatley, R Kumar, EJ Rodger, A Chatterjee, SD Diermeier, CJ Drummond, G Reid

## Abstract

Lung adenocarcinomas frequently harbour actionable oncogenic mutations that are vulnerable to treatment with targeted therapies. While responses to targeted therapies are often initially dramatic, relapse is almost inevitable and prevents durable responses in advanced-stage patients. Relapse is, in part, caused by drug tolerant persister cells (DTPs) which are able to survive treatment by entering a reversible, dormant state. Although long non-coding RNAs (lncRNAs) regulate processes thought to allow DTPs to survive and become stably resistant, the potential roles of lncRNAs in DTPs are largely unknown. In this study, we sought to investigate the expression of lncRNAs in *in vitro* DTP models of lung adenocarcinoma. We found that the lncRNAs *Metastasis-Associated Lung Adenocarcinoma Transcript 1* (*MALAT1*) and *Nuclear Paraspeckle Assembly Transcript 1* (*NEAT1*) were enriched in DTPs and that knocking down *MALAT1* enhanced the effect of targeted therapies in both EGFR- and KRAS-mutant DTP models. To better understand pathways that *MALAT1* might regulate in DTPs, bulk RNA-sequencing was performed and several pathways that may contribute to the actions of *MALAT1* in DTPs were identified. Overall, our work describes a role for the lncRNA *MALAT1* in DTPs in NSCLC and suggests that *MALAT1* may be a novel target for the prevention of drug tolerance and subsequent resistance to targeted therapy in NSCLC.

## Introduction

Advanced non-small cell lung cancer (NSCLC) is the most prevalent form of lung cancer^1–3^ and frequently harbours mutated oncogenes^4–7^. Although such tumours often respond well to targeted therapies^4,8^, the development of resistance is an almost inevitable outcome for advanced-stage patients^4^. It is now widely accepted that therapy resistance in NSCLC is, in part, mediated by drug tolerant persister cells (DTPs)^7,9,10^ that are largely defined by their ability to tolerate normally lethal drug doses in a non-genetic and reversible manner^9,11^. DTPs are then thought to acquire stable resistance to targeted therapies via “adaptive mutability”, a process proposed to generate *de novo* resistance-conferring mutations^12–14^. Despite having a heterogeneous phenotype, DTPs are broadly reliant on transcriptional re-programming as well as epigenetic and metabolic alterations to survive treatment^7,9,15–21^.

In acquiring these traits for survival, DTPs compromise other traits that usually confer fitness to the cell and thus provide potentially exploitable vulnerabilities^22,23^. For example, DTPs almost completely lose control over oxidation and reduction (redox) pathways by downregulating master redox regulators, such as nuclear factor erythroid 2-related factor 2 (NRF2)^24–26^. This leaves DTPs vulnerable to inhibition of the few intact reducing factors such as glutathione peroxidase 4 (GPX4)^26,27^, the cysteine–glutamate transporter system Xc^28^, and aldehyde dehydrogenase (ALDH)^29–31^. Another common strategy for targeting DTPs is by preventing or exploiting the epigenetic alterations that underpin the phenotype^9,11^. For example, DTPs are dependent on several histone demethylase events that are mediated by KDM5A^9^ and inhibition of KDM5A either directly using KDM5 inhibitors such as CPI-445^32^, or indirectly, by inhibiting upstream signalling, such as inhibition of IGF-1R^9^ has been found to inhibit DTPs. However, despite significant progress in our understanding of DTPs in NSCLC and the identification of exploitable vulnerabilities, none of the strategies designed to specifically target DTPs have been implemented clinically to date^11,33^. Without a clear clinical strategy to target DTPs, long-term responses to targeted therapies remains an elusive outcome for patients.

While metabolic^26,31^, epigenetic^9,34^ and protein-coding gene expression changes^7,35^ are now well understood in DTPs^11,23^, non-coding alterations, particularly changes to long non-coding RNAs (lncRNAs), are poorly characterised^36^. LncRNAs, which are arbitrarily defined as non-coding RNA transcripts longer than 200 nucleotides in length^37^, are known to contribute extensively to cancer biology^38,39^. Several lncRNAs, such as *MALAT1* and *NEAT1*, are known to regulate several processes thought to support DTPs^36^ and recent research has indicated that such lncRNAs, including *MALAT1* and *NEAT1*, are altered in this population^35^. Despite this, little effort has been made to investigate the potential contribution of lncRNAs to DTPs, leaving a gap in our understanding of the biology of this population and any vulnerabilities that may emerge as a result.

Here, we demonstrate that *MALAT1* and *NEAT1* are enriched in DTPs induced by therapies that target EGFR or KRAS^G12C^ in NSCLC. Knockdown of *MALAT1* further sensitises DTPs to treatment with targeted therapies and we identify several candidate pathways that could mediate this effect. Overall, this work presents a novel vulnerability in lung adenocarcinoma cells tolerant to targeted therapies in NSCLC.

## Methods

### Cell Lines and Culture

PC9 (EGFR^exon19del^, RRID:CVCL_B260) and H358 (KRAS^G12C^, RRID:CVCL_1559) lung adenocarcinoma cell lines were purchased from ATCC. HEK293T (RRID:CVCL_0063) cells were a gift from Dr Silke Neumann (University of Otago). PC9 and H358 cells were cultured in GlutaMAX RPMI-1640 (ThermoFisher Scientific), while HEK293T cells were cultured in DMEM (high glucose, pyruvate, ThermoFisher Scientific). Culture medium was supplemented with 10%v/v foetal bovine serum, and all cells were cultured in a humidified atmosphere at 37 °C in 5% CO_2_ in air. Cells were routinely tested for mycoplasma (MycoAlert, Lonza). Cell line verification was performed by Cellbank Australia. Both PC9 and H358 cells were considered a match by cell line verification.

### Molecular Cloning

The pSin-mCherry-PuroR plasmid was constructed using the pSin-PuroR backbone (from pSin-EF2-Nanog-PuroR, which was a gift from Dr James Thomson^40^, University of Wisconsin-Madison, Addgene #16578). pSin-PuroR was digested using SpeI-HF and EcoRI-HF (New England Biolabs) to remove Nanog. The mCherry insert was PCR amplified from pSIN-IRES-mCherry, which was a gift from Dr Mark Hatley (St Jude Children’s Research Hospital)^41^ using primers incorporating SpeI and EcoRI restriction sites (5’SpeI: TAAGCAACTAGTACACGATGATAAGCTTGCCAC, 3’EcoRI: TAAGCAGAATTCCTAGGTCTCGAGCGGCCATC) and blunt cloned into pCR™-Blunt II-TOPO (ThermoFisher). The TOPO-mCherry plasmid was then digested with SpeI-HF and EcoRI-HF, and purified by gel electrophoresis followed by extraction using the NucleoSpin Gel and PCR cleanup kit (Machery-Nagel). The mCherry amplicon was then ligated into pSin-PuroR using T4 DNA ligase (New England Biolabs). In-frame insertion was confirmed using Sanger sequencing (Genetic Analysis Services Otago, sequencing primers in **Supplementary Table 1**). pSin-mCherry-PuroR plasmid map is shown in **Supplementary Figure 1**.

### Viral Transduction

pSIN-IRES-mCherry^41^, along with pMDLg/pRRE, pRSV-Rev and pMD2.G (gifts from Didier Trono, University of Geneva, AddGene #2251, 12253, 12259)^42^ packaging and envelope plasmids, were co-transfected using Lipofectamine 3000 (ThermoFisher Scientific) into HEK-293T cell lines seeded at 5×10^5^/dish in 10 cm dishes. PC9 cells were transduced with virus-containing medium collected 48 hours post-transfection, following filtering using a 0.45 µM syringe.

### Drug Treatment

To generate DTPs, PC9 cells were treated with 2 μM osimertinib (SelleckChem S7297), and H358 cells were treated with 1 μM sotorasib (SelleckChem S8830). All drugs were stored in DMSO at −20 °C and were diluted further in culture medium immediately prior to use. Each experiment contained a DMSO solvent control, which was comparable to the highest DMSO content used in each experiment (%v/v).

### Antisense Oligonucleotide Knockdown

Antisense oligonucleotides were reverse-transfected at 20 nM into PC9 and H358 cells plated at 1×10^6^ cells/well in 10 cm culture dishes using Lipofectamine RNAiMAX (ThermoFisher Scientific). Sequences and modifications for *MALAT1*^43^ and *NEAT1*^44^ ASOs are found in **Supplementary Table 2**.

### RT-qPCR

RNA extraction was performed using the PureLink RNA Mini Kit (Ambion Life Technologies) according to the manufacturer’s protocol. Concentration and quality of RNA were determined via NanoPhotometer N60 nanodrop (Implen Inc). cDNA was produced using 1000 ng RNA and 5X qScript cDNA Supermix (Quantabio) and then diluted to 10 ng/µL for use. SYBR-based RT-qPCR was performed using the TB Green Premix Ex Taq II (Tli RNase H Plus) master mix kit (Takara) and 1 µL of 10 ng/µL cDNA. Primer sequences are available in **Supplementary Table 3**. RT-qPCR was performed using triplicate wells on QuantStudio’s Flex 7 qPCR machine and analysed following the 2^−ΔΔCT^ method^45^, with *ubiquitin C* (*UBC*) as a reference gene for normalisation.

### Proliferation Assays

PC9 and H358 cells were seeded at 1×10^3^ and 1.5×10^3^ cells/well respectively in 96-well plates and treated with respective targeted therapies 24 h following seeding. Following treatment for 5 days, culture plates were frozen at −80 °C. After thawing at room temperature, lysis buffer (20 mM Tris-HCL pH 8, 50 mM EDTA, 20% v/v Triton X100) containing 1:8000 SYBR Green I (ThermoFisher) was added and plates were incubated overnight in the dark at 4 °C. Plates were allowed to reach room temperature before being read at 485 nm excitation/535 nm emission on a Victor X4 plate reader (PerkinElmer). Data was normalised to relevant DMSO solvent controls and/or ASO-controls and graphed using GraphPad Prism 10 (Version 10.0.3).

### High-Content Microscopy

mCherry-PC9 cells were reverse transfected with 20 nM of relevant ASOs and plated at a density of 1×10^3^ cells/well in glass-bottomed 96-well culture plates (Griener). 2 μM osimertinib was added 18-24 hours following cell plating and was replaced weekly in fresh medium throughout the experiment. Every week, cells were imaged via Opera Phenix High Content Screening System (Revvity) in brightfield and mCherry channels using a 5× air objective. Images were acquired via Harmony (version 5.1, Revvity) software and analysed via Signals Image Artist (Version 1.0, Revvity). Cell confluency was calculated by the fraction of the well area occupied by cells. The total well area was determined using the brightfield channel, while the mCherry channel was used to measure areas occupied by cells.

### SplintR RT-qPCR

ASO concentration was quantified using the SplintR RT-qPCR assay^46^. In brief, extracted RNA was incubated with ligation probes and then heated for 5 minutes at 95 °C and cooled at a rate of 0.1 °C per second to 37 °C to hybridise the probes to MALAT1-AS1. Hybridised probes were then ligated using 25U/reaction of SplintR ligase (New England Biolabs) at 37 °C for 30 minutes. SplintR ligase was heat-inactivated at 65 °C for 20 minutes, and RT-qPCR was performed using a double-quenched TaqMan probe (FAM/ZEN/IABkFQ) to detect the ligated product. Sequences for SplintR probes and primers found in **Supplementary Table 4**.

### Homologous Recombination (HR) Assay

Homologous recombination capacity was quantified using the pDRGFP HR assay^47^. pDRGFP expressing PC9 and H358 cell lines were generated by transfecting 2.5×10^5^ cells/well with the pDRGFP plasmid^47^ (gift from Dr Maria Jasin, Memorial Sloan Kettering Cancer Center and Cornell University, AddGene #26475) using Lipofectamine 3000 (ThermoFisher). Puromycin (Gibco) was used to select stably transfected cells. Once generated, pDRGFP expressing cells were reverse co-transfected with ASOs, along with pCBASce-I^47,48^ (gift from Dr Maria Jasin, AddGene #26477) and pSin-mCherry-PuroR (**Supplementary Figure 1**) using FuGENE HD (Promega). Transfected cells were then treated with solvent control, 2 µM osimertinib (PC9) or 1 µM sotorasib (H358) for 72 hours before cells were harvested, stained with Zombie-NIR viability dye (1:1000, BioLegend) and analysed by flow cytometry using a BC CytoFLEX S flow cytometer (Beckman Coulter). Data was recorded using the CytExpert software (Beckman Coulter) and analysed using FloJo (BD Biosciences). Cells were gated as follows: acquisition and doublet exclusion, selection of viable cells (Zombie-NIR negative), selection of mCherry positive cells (control for transfection efficiency), selection of cells able to repair the pDRGFP plasmid (GFP). HR capacity was determined by normalising treated samples to the percentage of GFP-positive live and transfected (mCherry-positive) cells in treated samples to the ASO-control reverse-transfected, DMSO-treated controls. An average of three biological replicates was graphed using GraphPad Prism 10 software for MacOS (Version 10.0.3).

### RNA-Sequencing Analysis

RNA was extracted using the PureLink RNA Mini Kit (Ambion Life Technologies) according to the manufacturer’s protocol. rRNA depletion was performed, and RNA-seq libraries were constructed using the TruSeq Total RNA-Seq library prep kit (Illumina). Paired-end sequencing was performed on an Illumina NextSeq 2000 next-generation system (performed by Otago Genomics Facility, University of Otago).

RNA-Sequencing data assessment and analysis was performed using our previously described pipelines^49,50^. Briefly, Trim Galore! was used to perform adapter trimming and Spliced Transcripts Alignment to a Reference (STAR)^51^ was used to align cleaned sequenced reads to the reference human genome (GRCh37). Read counts were assigned to annotated genes using featureCounts^52^ with an average of 26.6 million reads per sample. Differential gene expression analysis was performed using DESeq2^53^. Gene ontology analysis was performed using clusterProfiler^54^ and was conducted using the org.Hs.eg.db annotation package. Gene ontology analysis was based on Biological Process (BP).

### Statistical Analysis

In RT-qPCR, HR and proliferation assays, an average of three biological replicates were analysed using unpaired one-tailed t-tests. High content imaging and flow cytometry experiments were analysed using Welch’s T-tests. A p-value < 0.05 was considered significant in these experiments. For RNA-sequencing investigating *MALAT1*-KD in DTPs, statistical analysis was performed using the BH method to generate adjusted p-values. For RNA-sequencing results, an adjusted p-value < 0.05 was deemed to be statistically significant.

## Results

### Drug Tolerant Persisters in EGFR- and KRAS-Mutant Lung Adenocarcinoma Models

To investigate lncRNAs in DTPs in lung adenocarcinoma, we first sought to develop *in vitro* models of both EGFR mutant and KRAS mutant DTPs. One of the defining features of drug-tolerant persisters (DTPs) is their ability to survive treatment despite high concentrations of targeted therapy^9,23^. To identify concentrations of targeted therapies which result in DTPs, we performed proliferation assays on the EGFR-mutant PC9 (EGFR^Del19^) treated with the third-generation EGFR inhibitor osimertinib and the KRAS-mutant H358 cell line (KRAS^G12C^) treated with the KRAS^G12C^ inhibitor sotorasib for 120 hours (5 days). As with previous studies^9,17^, we found that a small population of cells (approximately 10%) survived treatment with targeted therapies, even at doses of up to 50 µM **(Figure 1A and B)**.

**Figure 1:**
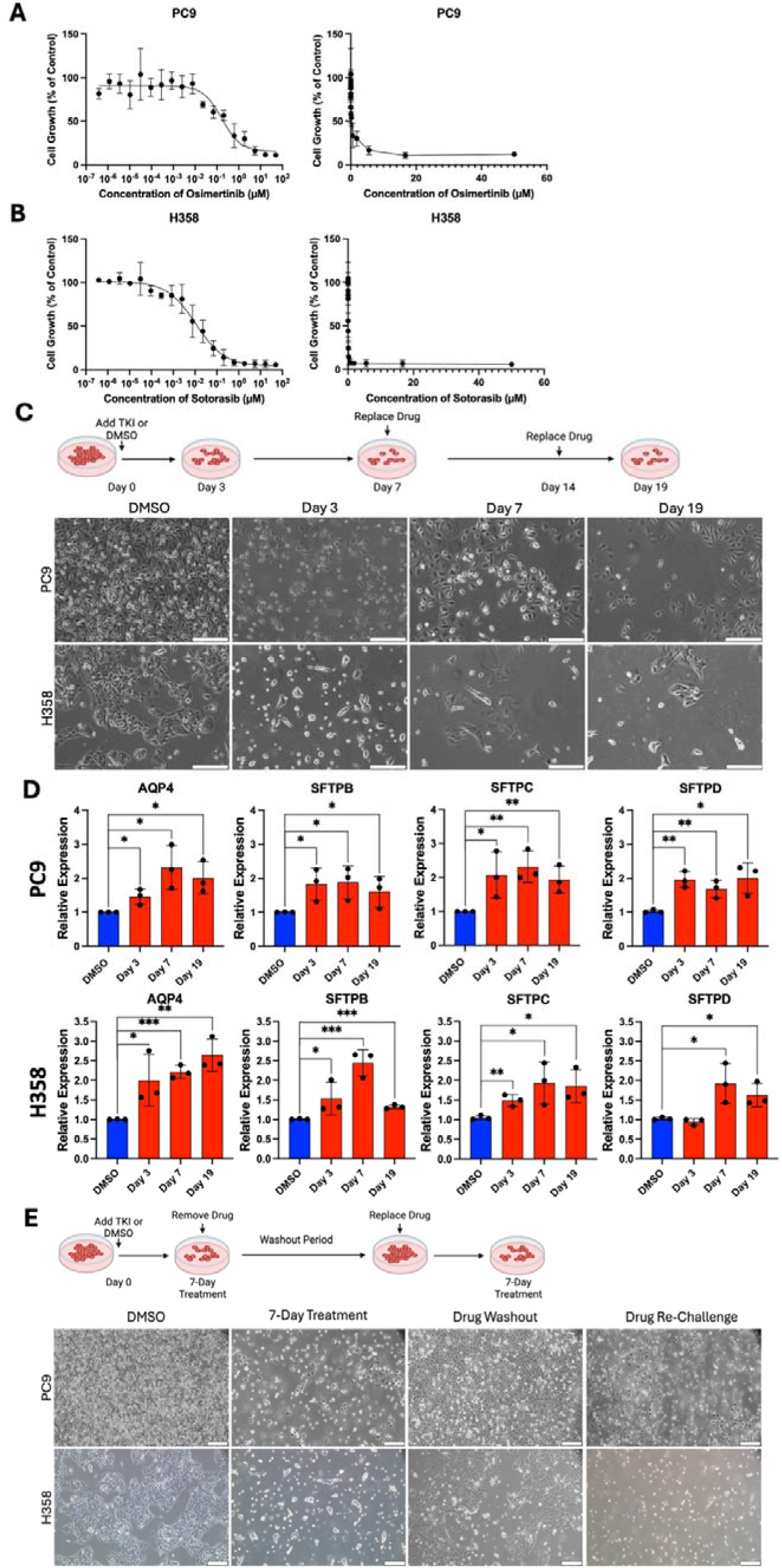
*In Vitro* Models of Drug Tolerant Persisters in NSCLC Cell Lines. **(A and B)** Proliferation assays were performed on PC9 cells **(A)** or H358 cells **(B)** treated with osimertinib or sotorasib respectively for 5 days. Each experiment was normalised to respective solvent controls, and data is displayed in log10 (left) and linear (right) formats. Data is mean ± standard deviation of 3 biologically independent replicates (n=3). **(C)** PC9 cells (top) or H358 cells (bottom) were treated with respective targeted therapy or DMSO %v/v for 3, 7, or 19 days. Cells were imaged via digital inverted brightfield microscopy. Representative images are shown (n=3). **(D)** Expression of minimal residual disease markers in DTPs by RT-qPCR. All genes are relative to a DMSO solvent control. Unpaired, one-tailed t-tests were performed to test for significance Data is mean ± standard deviation (n=3). *=p ≤ 0.05, **=p ≤ 0.01, ***=p ≤ 0.001, ****=p ≤ 0.0001. **(E)** Targeted therapy treatment, washout and re-challenge. Brightfield imaging of PC9 cells (top) and H358 cells (bottom) treated as indicated. Representative images are shown from 3 biologically independent repeats (n=3). Scale bar = 200 µm.

We then treated PC9 and H358 cell lines with osimertinib or sotorasib respectively, for 3, 7 or 19 days to generate DTP models. The treatment regimen (drug concentration and duration of exposure) was informed by proliferation assays **(Figure 1A and B)** and previous literature that compared residual disease in lung adenocarcinoma patients with *in vitro* models of drug tolerance^7^. Following treatment with targeted therapies, a small fraction of cells remained viable in both cell lines **(Figure 1C)**. Next, we investigated whether the expression of markers previously linked to DTPs and minimal residual disease in NSCLC patients^7,35^ were also expressed in our DTP models. PC9 cells demonstrated statistically significant increases in the expression of aquaporin 4 (*AQP4*) and the surfactants B, C and D (*SFTPB, SFTPC, SFTPD*)^7^ at all time points **(Figure 1D)**. H358 cells exhibited statistically significant increases in DTP markers *SFTPB, SFTPC* and *AQP4* at all time points and *SFTPD* after 7 and 19 days **(Figure 1D)**.

A key hallmark of drug tolerance is the reversible nature of the phenotype^9,55^. To further confirm that our DTP models behaved similarly to other models, we performed drug treatment, withdrawal and rechallenge experiments. We found that both PC9 and H358 cells rapidly proliferated upon drug withdrawal **(Figure 1E)**. Rechallenge with targeted therapy resulted in a similar response to initial drug treatment in both cell lines, as observed in other previously described DTP models^9,55^. When investigating expression of DTP markers, we found that markers fluctuated in expression upon drug treatment, washout and re-challenge in PC9 cells **(Supplementary Figure 2)**. Overall, these results confirm that the DTP models developed here are comparable to previous models of tolerance and behave similarly minimal residual disease in patients receiving treatment with targeted therapies^7,12,14,35^.

### EGFR and KRAS mutant DTP models Demonstrate Similar Changes in lncRNA Expression

Having reproduced *in vitro* models of DTPs in PC9 and H358 cells, we aimed to explore lncRNA expression in these models. To do so, we performed differential analysis using bulk RNA-sequencing on PC9 and H358 NSCLC cell lines treated with osimertinib or sotorasib compared to control (DMSO-treated), respectively for 3, 7, or 19 days. We identified several lncRNAs that were differentially expressed upon treatment and selected lncRNAs that demonstrated similar differential expression responses to treatment in both cell lines and were expressed at levels ≥ 1 TPM for further analysis **(Figure 2A)**. Using these criteria, we identified the lncRNAs *MALAT1, NEAT1, UFC1*, and *ANRIL* as potential candidates for further study, and confirmed these findings using RT-qPCR **(Figure 2E and F)**.

**Figure 2:**
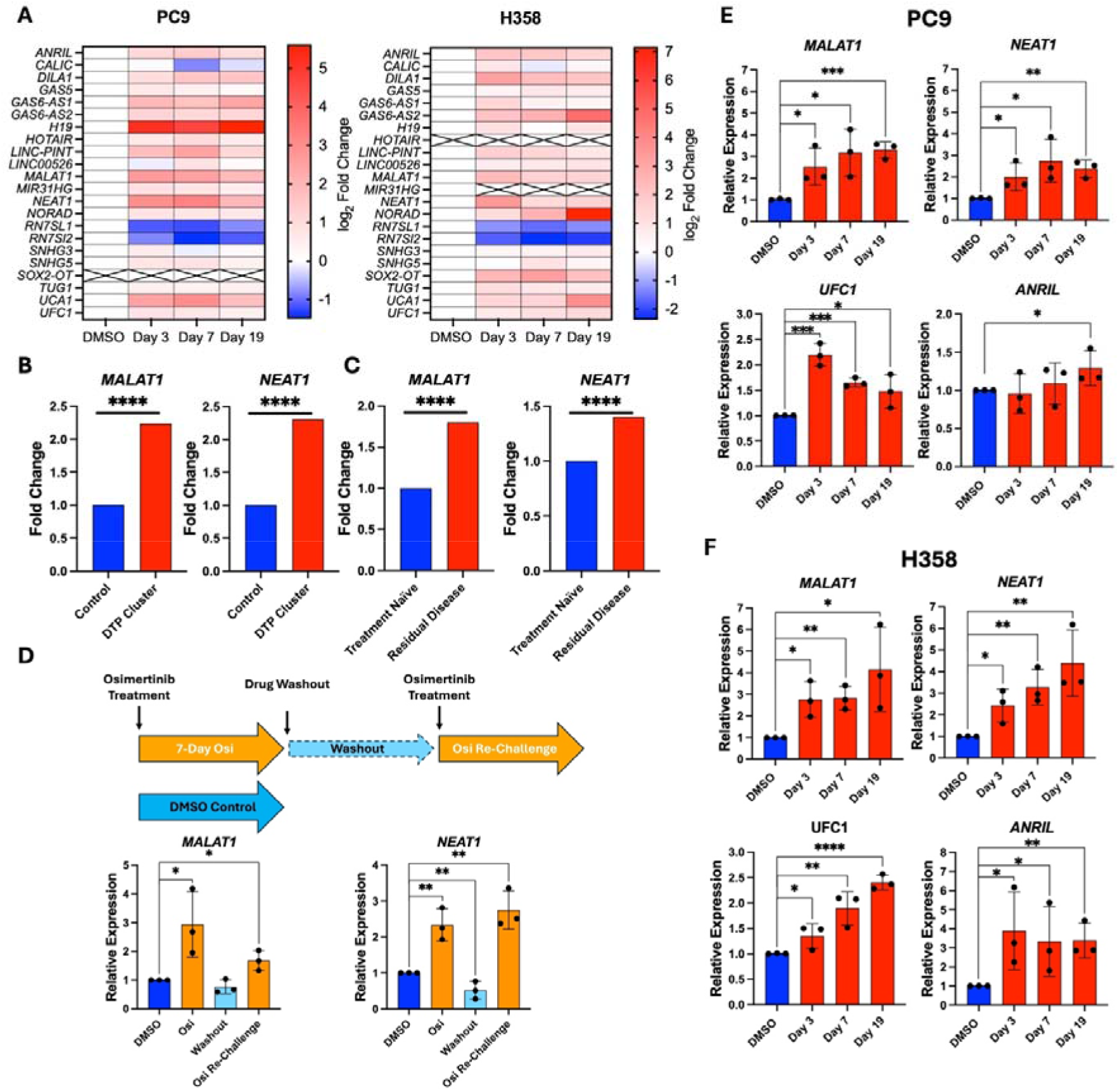
Expression of LncRNAs in Drug Tolerant Persisters in Lung Adenocarcinoma Models. **(A)** Expression of lncRNAs by RNA-seq analysis in PC9 and H358 cell lines treated with 2 µM osimertinib or 1 µM sotorasib respectively for 3, 7 and 19 Days. Heatmaps represent log2 fold change as calculated based on transcript per million data normalised to the DMSO control. Cells with crosses indicate that no reads were mapped to that transcript in the indicated condition. Data is from one biological repeat (n=1). **(B)** Expression of *MALAT1* and *NEAT1* in DTPs treated with erlotinib from public scRNA-seq data^35^. Differential expression analysis was performed by Benevolenskaya and colleagues^35^ using a DESeq2 approach. **** =adjusted p value ≤0.00001. **(C)** *MALAT1* and *NEAT1* expression in residual disease samples from lung cancer patients undergoing targeted therapy treatment. *MALAT1* and *NEAT1* expression was investigated in this dataset, based on differential expression analysis performed by the authors^7^. **** =adjusted p value ≤0.00001. **(D)** Relative expression of *MALAT1* and *NEAT1* in osimertinib treatment, washout and re-challenge experiments relative to solvent controls in PC9 cells. RNA was collected after: 7 days of treatment; 7 days of treatment followed by proliferation in drug-free medium; or 7 days of treatment followed by drug-free proliferation and then a 7-day re-challenge with targeted therapy. **(E and F)** RT-qPCR validation of lncRNA expression in PC9 **(E)** and H358 **(F)** DTPs. All genes are relative to a DMSO solvent control. Unpaired, one-tailed t-tests were performed to test for significance. *=p ≤ 0.05, **=p ≤ 0.01, ***=p ≤ 0.001, ****=p ≤0.0001. Data is mean ± standard deviation (n=3).

To identify lncRNAs that overlap between our study and previous literature, we investigated publicly available single cell RNA-seq (scRNA-seq) data on PC9 DTP models treated with 2 μM of the EGFR-inhibitor erlotinib (GEO accession number GSE148465)^35^. A total of 11 lncRNAs were identified as differentially expressed across cells within DTP clusters with statistical significance (adjusted p-value <0.05) in gene expression data generated by Aissa and colleagues. *MALAT1* and *NEAT1* were the most strongly upregulated lncRNAs among drug treated DTP clusters **(Figure 2B)**, showing 2.2-fold and 2.3-fold increases in expression respectively compared to solvent (DMSO) controls, reflecting our findings.

Next, we assessed whether enrichment of *MALAT1* and *NEAT1* was an *in vitro* specific effect or whether this phenomenon would extend to lung cancer patients receiving treatment with targeted therapies. To address this, we investigated a scRNA-seq dataset^7^ derived from patient samples (49 clinical biopsies obtained from 30 patients before and during targeted therapy treatment, NCBI BioProject #PRJNA591860) that investigated the molecular features of NSCLC at targeted therapy naïve and residual disease stages. When comparing residual disease to targeted therapy naïve sample data generated by the authors, *MALAT1* and *NEAT1* were enriched by 1.8- and 1.4-fold respectively (adjusted p-value <0.05, **Figure 2C)**.

To investigate *MALAT1* and *NEAT1* further in a DTP context, we performed drug treatment, washout and re-challenge assays **(Figure 2D)**. We found that increases in both *MALAT1* and *NEAT1* expression was transient upon drug treatment and reversible with drug withdrawal. After cells were exposed to re-treatment, *MALAT1* and *NEAT1* expression once again increased (1.7- and 2.7-fold respectively compared to DMSO controls following drug re-challenge).

### MALAT1 Contributes to Drug Response and DTP Survival

To investigate the roles of *MALAT1* and *NEAT1* in drug tolerance, we performed antisense oligonucleotide (ASO)-mediated knockdown of *MALAT1* and *NEAT1*. We began by testing the knockdown ability of ASOs designed to target *MALAT1* and *NEAT1* at a concentration of 20 nM in PC9 and H358 cells **(Figure 3A and B)**. Each of the two ASOs designed to target *MALAT1* significantly reduced the abundance of the transcript in PC9 and H358 cell lines 48 hours post-transfection (an approximate 90% and 80% decrease, respectively, compared to control ASO). Likewise, all four *NEAT1*-targeting ASOs significantly reduced the abundance of *NEAT1* in both cell lines after 48 hours (an approximate 80% decrease in both cell lines compared to control ASO). No obvious changes to morphology were observed when knocking down either *MALAT1* or *NEAT1* **(Figure 3A and B, left)**. For subsequent experiments, one ASO for each lncRNA was selected (MALAT1-AS1 and NEAT1-AS1) as these ASOs elicited the strongest knockdown effect on each cell line tested. *MALAT1* and *NEAT1* are known to be involved in the regulation of similar subsets of genes^56,57^. Because of this, it was essential to investigate whether knockdown of one lncRNA changes the expression or enrichment level of the other. We found that knockdown of *MALAT1* or *NEAT1* with corresponding ASO did not alter the abundance of the reciprocal gene **(Supplementary Figure 3A and B)**.

**Figure 3:**
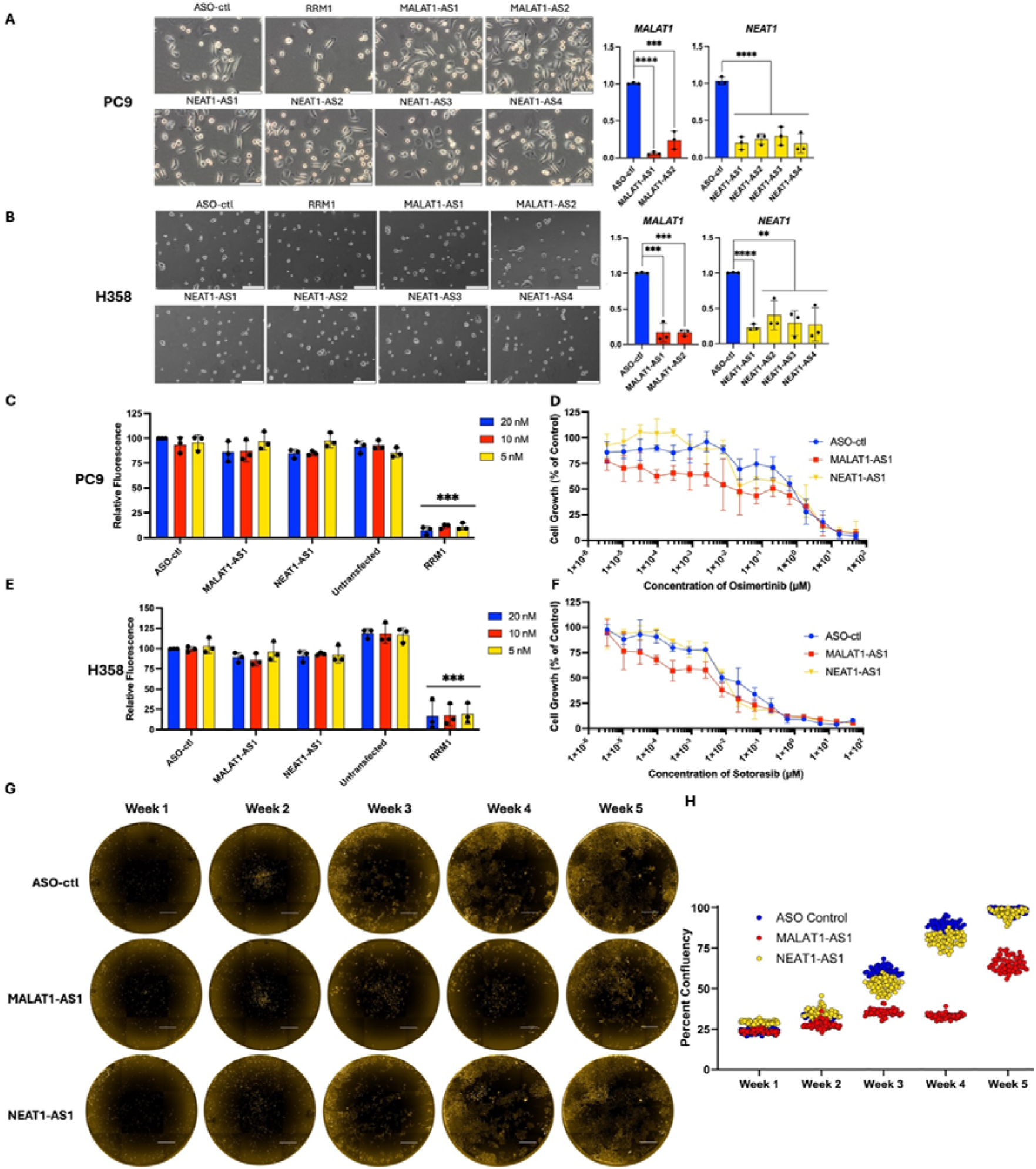
*MALAT1* Contributes to Targeted Therapy Response and DTP Survival. **(A and B)** ASO-mediated knockdown of *MALAT1* and *NEAT1* in PC9 **(A)** and H358 **(B)** cells. Left: brightfield images of cells post-transfection at indicated time points with ASOs indicated. Scale bar = 200 µm. Right: RT-qPCR for *MALAT1* and *NEAT1*. All genes are relative to a DMSO solvent control. Data is mean ± standard deviation (n=3). Unpaired, one-tailed t-tests were performed to test for significance. Significance *=p ≤ 0.05, **=p ≤ 0.01, ***=p ≤ 0.001, ****=p ≤ 0.0001. **(C-F)** Proliferation assays were performed on PC9 cells **(C and D)** or H358 cells **(E and F)** either with multiple concentrations of ASO (**C and E**) or with relevant targeted therapy (**D and F**). 72 hours after transfection, T-tests were performed by unpaired one-tailed t-test. Significance: *=p ≤ 0.05, **=p ≤ 0.01, ***=p ≤ 0.001. Data is mean ± standard deviation (n=3). **(G)** High-content imaging of long-term osimertinib treatment in combination with *MALAT1* or *NEAT1* knockdown in PC9-mCherry cells. Cells were reverse transfected with either a control ASO, MALAT1-AS1, or NEAT1-AS1 and treated with osimertinib for 5 weeks. Each week, cells were imaged in the mCherry channel. Images were processed using Harmony (version 5.1, Revvity). Images shown are representative images obtained from 180 wells per condition. Scale bars = 1 mm. **(H)** Analysis of high-content imaging data. Blue = ASO-control, red = *MALAT1*-KD, yellow = *NEAT1*-KD. Confluency analysis was performed in Signals Image Artist (Revvity) software and graphed in GraphPad Prism.

We then investigated cell viability following *MALAT1* or *NEAT1* knockdown in our lung adenocarcinoma cell lines by performing proliferation assays. In both cell lines, no significant growth inhibition was observed in cells transfected with 5 nM, 10 nM or 20 nM of MALAT1-AS1 or NEAT1-AS1 compared to those transfected with 20 nM of ASO-control **(Figure 3C and E)**. The positive transfection control, siRRM1, which is known to strongly inhibit the ribonucleotide reductase M1 subunit (*RRM1*)^58^, significantly reduced cell viability in both cell line models, indicating successful transfection in both conditions **(Supplementary Figure 3C)**.

We hypothesised that knocking down *MALAT1* or *NEAT1* could improve the efficacy of targeted therapies in lung adenocarcinoma cell lines. To test this, we reverse transfected 96-well plates with either an ASO-control oligonucleotide, MALAT1-AS1 or NEAT1-AS1 at a 20 nM concentration 18-24 hours prior to treatment with targeted therapies. We found that knockdown of *MALAT1* enhanced the response of cells to targeted therapies following 72 hours of exposure in both PC9 and H358 cell lines **(Figure 3D and F)**. In PC9 cells, this effect response was greatest between 94 pM and 0.2 µM of osimertinib, while in H358 cells this response was greatest between 94 pM and 7.62 nM of sotorasib. In contrast, knockdown of *NEAT1* had no significant effect in either cell line.

To investigate the effect of *MALAT1* and *NEAT1*-KD in DTPs at longer time points we performed high-content imaging over a period of five weeks using PC9 cells constitutively expressing mCherry to enable quantification of cell confluency. Proliferation assays were used to confirm negligible effects on cell growth from constitutive expression of mCherry **(Supplementary Figure 3D)**. When examining the mCherry fluorescent channel of the images generated from high-content imaging, it became apparent that *MALAT*-KD enhanced the inhibitory effect of osimertinib treatment in PC9 cells **(Figure 3G)**. When measuring cell confluency, we found a significant difference in cell confluency between *MALAT1*-KD and ASO-control transfected DTPs under osimertinib treatment **(Figure 3H)**. This finding was most apparent between weeks 3 and 4 of treatment. At week 3, the mean confluency was approximately 30% in the *MALAT1*-KD group compared to 60% and 50% in the control and *NEAT1*-KD groups respectively. At week 4, mean confluency was 30% in the *MALAT1*-KD group compared to 90% in the control group and 80% in the *NEAT1*-KD group. At week 5, the difference in confluency between *MALAT1*-KD and control transfections began to diminish, with the *MALAT1*-KD group was 65% confluent on average, compared to 98% and 96% average confluency for control and NEAT1-AS1 groups respectively. We hypothesized that the effect of MALAT1-AS1 enhanced the effect of targeted therapy and that effect diminished over time due to degradation or dilution of the ASO. To test this, we quantified the abundance of the ASO using the SplintR qPCR assay^46^ and found that the effect size correlated with a the presence of MALAT1-AS1 in DTPs and had an inverse relationship with expression of *MALAT1* **(Supplementary Figure 3E and F)**.

### *MALAT1*-KD Does Not Further Alter Targeted Therapy-Mediated Changes in Adaptive Mutability in DTPs

DTPs have been demonstrated to act as a pool^59^ of cells in which resistance-conferring mutations may occur, leading the cancer to become genetically resistant to the targeted therapy via “adaptive mutability^11,12,23^”. As *MALAT1* has previously been linked to DNA damage responses^60,61^, we next investigated markers of adaptive mutability^12,14^, including markers for DNA mismatch repair (MMR), homologous recombination (HR) and DNA polymerase activity **(Figure 4A and B)**. *MALAT1* has been previously linked to the regulation of HR in untreated NSCLC cell models via regulation of *BRCA1*^60–62^. We first investigated adaptive mutability in our models of DTPS in the absence of ASOs and found several genes involved in MMR (*MLH1* and *MSH6*), HR (*BRCA1* and *BRCA2*) and canonical (high-fidelity) polymerase (*POLE, POLD* and *POLA*) pathways to be downregulated in both PC9 and H358 treated with osimertinib or sotorasib respectively for 3, 7 and 19 days. Additionally, the translesion polymerase, *POLI*, was upregulated by treatment at all time points in H358 DTPs and was upregulated in response to treatment at days 7 and 19, but not at day 3 in PC9 DTPs.

**Figure 4:**
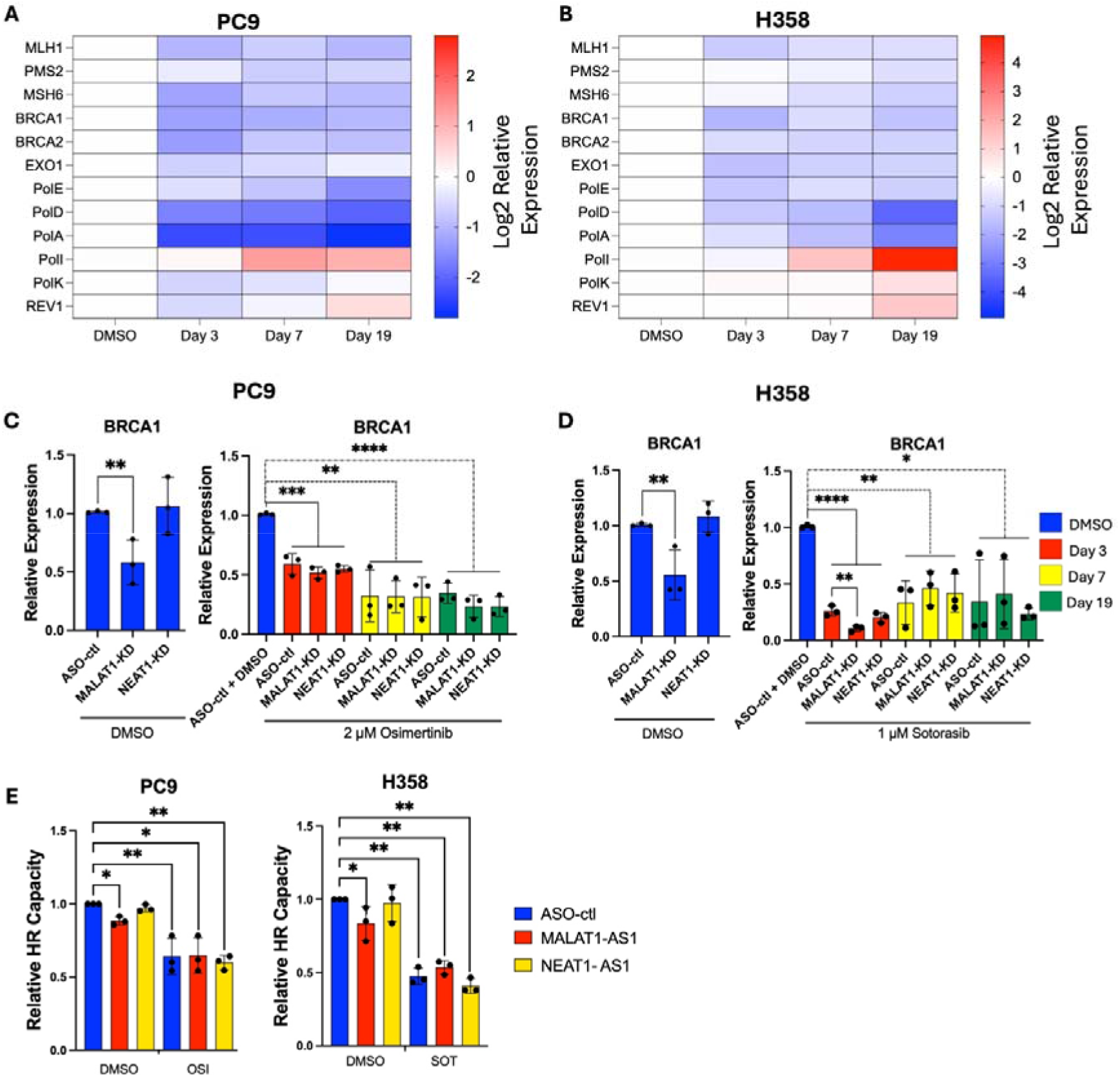
Knockdown of *MALAT1* Does Not Further Alter Targeted Therapy-Mediated Changes in Adaptive Mutability in DTPs. **(A and B)** Expression of adaptive mutability associated genes in PC9 cells treated with osimertinib **(A)** or H358 cells treated with sotorasib **(B)** by RT-qPCR. All genes are relative to a DMSO solvent control. Unpaired, one-tailed t-tests were performed to test for significance. *=p ≤ 0.05, **=p ≤ 0.01, ***=p ≤ 0.001, ****=p ≤ 0.0001. Data is mean ± standard deviation (n=3). **(C and D)** Relative expression of *BRCA1* and *BRCA2* in PC9 **(C)** or H358 **(D)** DTPs +/−*MALAT1* or *NEAT1* knockdown. Left: cells were transfected with either an ASO-control, MALAT1-AS1 or NEAT1-AS1 and treated with a %v/v DMSO solvent control for 3 days. Right: cells were transfected with either an ASO-control, MALAT1-AS1 or NEAT1-AS1 and treated with osimertinib or sotorasib for 3 days, 7 days or 19 days or with a DMSO solvent control for 3 days. All genes are relative to a DMSO solvent control. *=p ≤ 0.05, **=p ≤ 0.01, ***=p ≤ 0.001, ****=p ≤ 0.0001. Data is mean ± standard deviation (n=3). (**E)** Relative HR capacity was determined by normalising the percentage of GFP expressing cells to the GFP signal in ASO-control, DMSO treated sample. Plots were generated via FlowJo (v10.10). T-tests were performed by unpaired one-tailed t-test. ns = not significant *=p ≤ 0.05, **=p ≤ 0.001, Data for each experiment is mean ± standard deviation (n=3).

To investigate whether *MALAT1* enhances the survival of DTPs under treatment via adaptive mutability, we knocked down *MALAT1* or *NEAT1* in PC9 and H358 cells before treating them with either osimertinib or sotorasib respectively for 3, 7, or 19 days. We found that knockdown of *MALAT1* reduced *BRCA1* expression by approximately 50% in comparison to control transfections in both cell lines **(Figure 4C and D)**. Osimertinib and sotorasib treatment reduced expression of BRCA1 by approximately 50% and 70% respectively compared to DMSO solvent controls **(Figure 4C and D)**. Despite this, MALAT1 did not further reduce *BRCA1* expression down in targeted therapy treated cells at any time point, except for a modest decrease at 72 hours of treatment in H358 cells **(Figure 4C and D)**. *NEAT1*-KD did not change expression of *BRCA1* in either cell line with or without the addition of targeted therapy. *MALAT1* and *NEAT1* were knocked down effectively in our DTP models over the time points and in the conditions investigated **(Supplementary Figure 4A and B)**.

We then performed a plasmid-based HR assay^12,47^ to investigate whether knockdown of *MALAT1* or *NEAT1* has functional implications on HR capacity in DTPs^12,47^. In both PC9 and H358 cells, treatment with osimertinib or sotorasib respectively reduced the capacity of HR by 45-50%, matching previous reports in osimertinib treated PC9 cell lines^14^ **(Figure 4E and Supplementary Figure 5A and B)**. Knockdown of *MALAT1* consistently reduced the capacity of HR by 11% and 25% respectively in solvent-control (DMSO) PC9 and H358 samples. However, this effect was not additive upon treatment with targeted therapies in either cell line. Further, we found no statistical difference in either cell lines when comparing *NEAT1*-KD cells to ASO-controls in solvent control or targeted therapy treated cells.

### Knockdown of *MALAT1* Alters Expression of Pathways Related to the Drug Tolerant Persister Phenotype

To investigate potential mechanisms of action behind *MALAT1*-KD sensitising DTPs to targeted therapies, we performed bulk RNA-sequencing on samples obtained from PC9 cells +/−*MALAT1*-KD and treated with osimertinib for 3, 7, or 19 days. We found 200 genes were differentially expressed across each treated *MALAT1*-KD group compared to untransfected treated groups at 3, 7 and 19 days of osimertinib treatment (**Figure 5A and B**, adjusted p-value <0.05 and |log□FC| > 1, **Supplementary Table 5**).To investigate pathway changes elicited by *MALAT1*-KD in DTPs, we divided differentially expressed genes that were shared between time points into upregulated and downregulated groups and performed gene ontology (GO) analysis using clusterProfiler^54^ **(Figure 5C)**. We found that ontologies related to the inflammatory responses (GO:0050727, GO:0002526) were highly enriched by *MALAT1*-KD, while processes involved in Wnt-signalling were downregulated following *MALAT1*-KD. Several IL-6 cytokine family members (IL-6, IL-11, IL-27, LIF, OSM), as well as several genes (SNAI1, CDH2, MMP2, MMP9) involved in epithelial-mesenchymal transition and inhibition of plasminogen activation signalling (*SERPINE1, SERPINB2, PLAU, PLAT, PLAUR*) were differentially expressed following *MALAT1*-KD in DTPs across all time points tested **(Figure 5D)**. *MALAT1*-KD appears to augment therapy-induced increases in *IL-6* and *SERPINE1* expression **(Figure 5E)**. To confirm our bulk RNA-sequencing finding, we used an orthogonal RT-qPCR approach and observed highly significant upregulation of *SERPINE1* and *IL-6* expression in drug treated cells across all time points (day 3, 7 and 19, **(Figure 5F and G)**.

**Figure 5:**
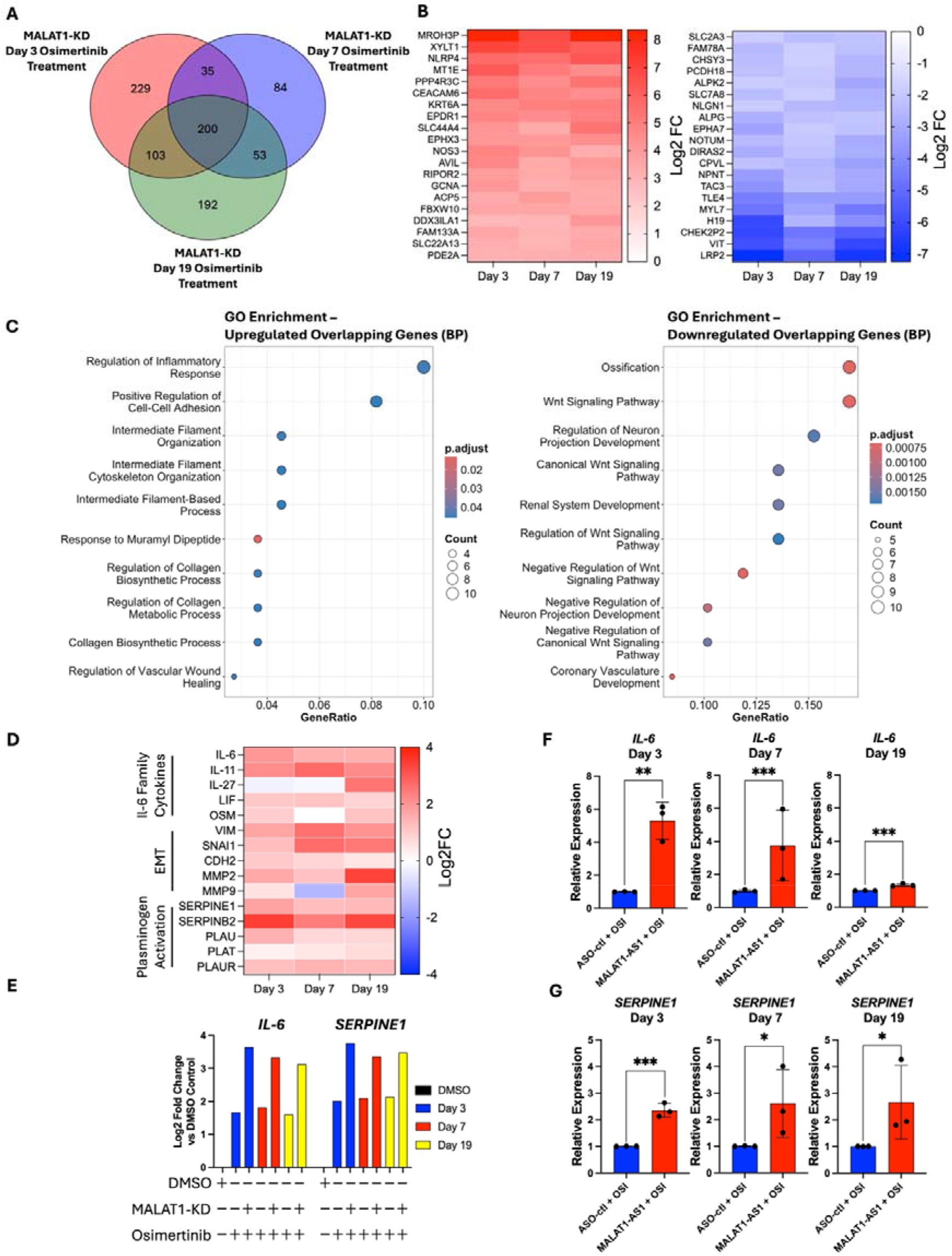
Knockdown of *MALAT1* Alters Expression of Pathways Related to the DTP Phenotype. **(A)** Venn diagram of concordant overlapping differentially expressed genes differentially expressed in *MALAT1*-KD PC9 DTPs compared to untransfected PC9 DTPs. **(B)** Heatmaps of top 20 concordant and overlapping genes altered by *MALAT1*-KD between treatment periods by log2 fold-change. Intensity represents log2 fold change. Red = log2 fold change > +1. Blue = log2 fold change < −1. All genes shown have adjusted p-values < 0.05. **(C)** Gene ontology analysis of the top 15 upregulated and downregulated pathways on genes differentially expressed by *MALAT1*-KD between 3, 7 and 19 days of osimertinib treatment. Circle shading reflects adjusted p-value, and circle circumference indicates the number of genes contributing to each biological process. GO analysis was performed using the clusterProfiler^54^ package in R Studio. **(D)** Gene expression of members the IL-6 cytokine family, EMT pathway and plasminogen activation signalling pathways by RNA-seq. **(E and F)** Expression of *IL-6* **(E)** and *SERPINE1* **(F)** by RT-qPCR in *MALAT1*-KD + osimertinib samples relative to ASO-control plus osimertinib samples at 3, 7 or 19 days as indicated. T-tests for RT-qPCR were performed by unpaired one-tailed t-test. Significance: *= p ≤ 0.05, **=p ≤ 0.01, ***=p ≤ 0.0001 Data is mean ± standard deviation (n=3)

## Discussion

Treatment with targeted therapies offers life-extending therapy for individuals diagnosed with advanced-stage NSCLC^63,64^. Targeted therapies offer patients longer progression-free survival compared with more traditional systemic approaches such as chemotherapies^63,65–68^. Despite initial clinical responses, targeted therapies are never curative in advanced-stage disease, with drug resistance and disease progression being inevitable^23^. Preventing non-genetic adaptation to targeted therapies, such as drug tolerance, remains an unsolved problem that leads to relapse in patients^11^. While considerable efforts have been mounted to address the issue of drug tolerance in NSCLC, no strategies to eliminate DTPs in a clinical setting have been successful to date^11,23^. In this study we aimed to investigate the lncRNAs *MALAT1* and *NEAT1* in drug tolerance in NSCLC.

Having established models of EGFR and KRAS mutant DTPs^7,9,35^, we demonstrated that the lncRNAs *MALAT1* and *NEAT1* are highly enriched in both models. Enrichment of *MALAT1* and *NEAT1* has been previously reported in erlotinib-induced DTPs^35^; however, their function and whether they are similarly enriched in other DTP models has not been investigated to date. We extend this data by investigating the role of *MALAT1* and *NEAT1* in osimertinib-treated DTPs in an EGFR-mutant cell line, and sotorasib-treated DTPs in a KRAS mutant cell line over extended treatment periods. Furthermore, by investigating *in vivo* data generated by Maynard *et. al*.^7^ our work reveals that this phenomenon may extend to patients treated with targeted therapies. Several factors might be inducing enrichment of *MALAT1* and *NEAT1* in DTPs, for example, DTPs are largely in G1 arrest^9,17^ and expression of *MALAT1* is thought to increase during the G1 phase of the cell cycle^43^. Expression of both *MALAT1* and *NEAT1* have also been linked to DNA damage^69–71^, which has been consistently shown to accumulate in a tolerant state^11,14,72^ and thus, could also explain this finding.

We discovered that knockdown of *MALAT1* using ASOs delays the emergence of DTPs in NSCLC cell lines treated with EGFR- and KRAS-targeted therapies. While there is limited research reporting the role of *MALAT1* in DTPs, previous work has described a relationship between resistance to erlotinib, a first-generation EGFR-inhibitor, and *MALAT1*^73^. This work described an increase in IC_50_ in *MALAT1* overexpression models, and a reciprocal decrease in cell viability at 72 hours when *MALAT1* was knocked down using siRNAs. The mechanism of action proposed in this study involved MALAT1 acting as a competing endogenous RNA for miR-125, which is thought to inhibit Rab25^73^, a GTPase linked to inhibition of apoptosis and cell proliferation^74^.

*MALAT1* is often highly enriched in NSCLC compared to healthy lung tissues^75,76^, and thus the biology of *MALAT1* is generally well understood in a pre-treatment NSCLC context^77^. *MALAT1* is often cited as a robust marker for poor prognosis^75,77,78^ and appears to regulate metastatic phenotypes in NSCLC^78,79^. Interestingly, *MALAT1* has also been linked to the reactivation of dormant cells following treatment with immunotherapies in breast cancers^56^, and several studies have demonstrated that knockdown of *MALAT1* further sensitises NSCLC cells to chemotherapies such as cisplatin^60,80^. Together these results could point to *MALAT1* having a role in pan-treatment tolerance and resistance that may not be specific to targeted therapies.

A key feature of DTPs is their ability to eventually acquire de novo resistance-conferring mutations, while in a predominantly slow-cycling state^11,12,81^. Adaptive mutability is primarily thought to be regulated via changes elicited through alterations in mRNA abundance, leading to dysregulation in DNA damage repair processes and heightened mutational rates in DTPs^12,14^. *MALAT1* is often observed to be upregulated upon induction of DNA damage^69^ and is well known to be a regulator of HR by regulating BRCA1^60,61^ in untreated NSCLC cell models^60^. In this role, inhibition of *MALAT1* has been proposed as a method of increasing sensitivity to PARP inhibitors in various cancer models^60–62^. However, despite regulating HR at the transcriptional level in solvent controls, we found that this effect was masked by targeted therapy treatment and thus is unlikely to contribute to the phenotypic effect of *MALAT1* expression in drug tolerance.

Several previous studies have investigated *MALAT1*-KD in a NSCLC context that draw parallels to our study, for example, Diederichs et. al. found that genetic knockout of MALAT1 in untreated A549 lung adenocarcinoma cell lines downregulated several EMT and metastasis genes^79^. Furthermore, a reduction in metastatic potential was observed when mice were implanted with *MALAT1*-KO human NSCLC cell lines^79^. In breast cancer models, *MALAT1* also appears to contribute to metastasis^82,83^ via TEAD related signalling^83^ and regulating alternative splicing^82^. Overall, we found similar differential expression patterns to these studies, particularly when comparing the GO analysis following *MALAT1*-KD between studies. We identified differential expression of IL-6 and plasminogen activation signalling in *MALAT1*-KD DTPs vs control transfected DTPs. Both IL-6 and plasminogen activation signalling are well known to be active during drug tolerance and have previously been linked to the DTP phenotype^7,35,84–86^, as well as mechanisms of resistance to targeted therapy^87–89^. Furthermore, related pathways, such as the JAK/STAT pathway, EMT, and SASP pathways, have also been linked to the DTP phenotype^84,87,90^. In the present study, we demonstrate an increase in expression of *IL-6* and *SERPINE1* upon treatment with targeted therapies, which is further increased upon knockdown of *MALAT1*. There is relatively limited literature linking *MALAT1* to IL-6. This literature is often contradictory and may be dependent on cell line and growth conditions. For example, knockdown of *MALAT1* in leukaemia, human umbilical vein endothelial (HUVEC) and breast cancer xenograft models in mice significantly increased expression of *IL-6*^56,91,92^. Furthermore, ASO-mediated knockdown in osteosarcoma and liver cancer cell lines elicits a strong upregulation of *IL-6* (between six and seven-fold)^43^. However, knockdown of *MALAT1* in gastric cancer cell lines results in the reciprocal effect, and IL-6 expression appears to be reduced by *MALAT1*-KD in such models^93^. Correlation studies investigating *MALAT1* expression in subtypes of breast cancer have found a strong negative correlation between expression of *MALAT1* and *SERPINE1*; however this appears to be subtype-specific at least in breast cancers^94^, and somewhat contradicts other literature that claims that *MALAT1* positively regulates other members of the Serpin family^56^.

There could be several scenarios that explain this, for example, expression of *IL-6* and *SERPINE1* are linked to cell stress response pathways^88,95^. It is therefore plausible that toxicity induced by targeted therapies upregulates these factors, and when toxicity is further augmented by *MALAT1*-KD, these factors further increase in expression as a result of increased cell toxicity. Interestingly, knockdown or knockout of *SERPINE1* has previously been performed in DTPs in NSCLC, eliciting a slight but statistical difference in DTP survival under treatment at longer (9 and 21 day) time points^35^. Another possibility is that *MALAT1* acts as a buffer for *IL-6* and *SERPINE1* to prevent overactivation of these pathways. For example, while overexpression of IL-6 often promotes proliferation^88^, high levels of IL-6 are known to induce cell dormancy and senescence^84,96,97^ and can lead to overactivation of p21 signalling^98–100^, which together could lead to cell cycle arrest in proliferative persisters^17,84^ and/or increase cell death. It is also likely that MALAT1-KD is elicits pleotropic effects^77^ in DTPs, impacting a number of process that may contribute to survival of DTPs.

While we observed a significant reduction in DTP viability under treatment following *MALAT1*-KD, a considerable amount of further work is required before considering any clinical applications of these findings. The first important question is the specific mechanism by which *MALAT1* enhances the effect of treatment in DTPs. Future studies in pre-clinical animal models will also be useful in ascertaining whether *MALAT1*-KD in combination with targeted therapies is a strategy to delay recurrence; in mice, *malat1*^−/−^ is not fatal and does not appear to confer phenotypic or histologic abnormalities^101,102^. While knockdown of *MALAT1* in human tissue cultures remain viable^101^, the long-term implications for inhibition of *MALAT1* in humans are unknown.

While ASO-based approaches have traditionally faced challenges in relation to stability and delivery^103,104^, recent advancements modifications that confer stability to ASOs, such as phosphothioate-linkages, and 2’methoxyethyl RNA modifications^103,105^, as well as delivery methods such as selective organ targeting (SORT) delivery methods that use lipid nanoparticle (LNP) delivery rapidly advanced to deliver stable oligonucleotides with specificity to a target organ, including the lung^106,107^. Such advancements have led to several recent clinical trials^108^ and the recent FDA approval of the ASO Tofersen for the treatment of SOD1-positive amyotrophic lateral sclerosis (ALS)^109^. An alternative approach to target *MALAT1* clinically could be the use of small molecules. Due to the uniquely stable 3’-expression and nuclear retention element (3’-ENE) structure of *MALAT1*^110^, small molecules have been developed to target *MALAT1*, which appear to be broadly successful in pre-clinical models^111–113^. Furthermore, re-purposed small molecule drugs, such as the well-tolerated and FDA-approved anthelmintic drug niclosamide^114,115^, have been shown to inhibit *MALAT1*^116^, meaning that a re-purposing strategy using small molecules to inhibit *MALAT1* in DTPs could be possible.

In conclusion, we find that the lncRNAs *MALAT1* and *NEAT1* are enriched in *in vitro* DTP models in NSCLC and that knockdown of *MALAT1* reduces DTP viability under treatment with targeted therapies. The results presented in this study suggest that inhibition of *MALAT1* may be an exploitable vulnerability for DTPs in lung adenocarcinoma.

## References

1 Molina, J. R., Yang, P., Cassivi, S. D., Schild, S. E. & Adjei, A. A. Non-small cell lung cancer: epidemiology, risk factors, treatment, and survivorship. Mayo Clin Proc 83, 584–594 (2008). 10.4065/83.5.584

2 Chen, Z., Fillmore, C. M., Hammerman, P. S., Kim, C. F. & Wong, K.-K. Non-small-cell lung cancers: a heterogeneous set of diseases. Nat Rev Cancer 14, 535–546 (2014). 10.1038/nrc3775

3 Herbst, R. S., Morgensztern, D. & Boshoff, C. The biology and management of non-small cell lung cancer. Nature 553, 446–454 (2018). 10.1038/nature25183

4 Rotow, J. & Bivona, T. G. Understanding and targeting resistance mechanisms in NSCLC. Nat Rev Cancer 17, 637–658 (2017). 10.1038/nrc.2017.84

5 Cancer Genome Atlas Research Network. Comprehensive molecular profiling of lung adenocarcinoma. Nature 511, 543–550 (2014). 10.1038/nature13385

6 Rosell, R. et al. Screening for epidermal growth factor receptor mutations in lung cancer. N Engl J Med 361, 958–967 (2009). 10.1056/NEJMoa0904554

7 Maynard, A. et al. Therapy-Induced Evolution of Human Lung Cancer Revealed by Single-Cell RNA Sequencing. Cell 182, 1232–1251.e1222 (2020). 10.1016/j.cell.2020.07.017

8 Lee, Y. T., Tan, Y. J. & Oon, C. E. Molecular targeted therapy: Treating cancer with specificity. Eur J Pharmacol 834, 188–196 (2018). 10.1016/j.ejphar.2018.07.034

9 Sharma, S. V. et al. A chromatin-mediated reversible drug-tolerant state in cancer cell subpopulations. Cell 141, 69–80 (2010). 10.1016/j.cell.2010.02.027

10 Haderk, F. et al. Focal adhesion kinase-YAP signaling axis drives drug-tolerant persister cells and residual disease in lung cancer. Nature Communications 15, 3741 (2024). 10.1038/s41467-024-47423-0

11 Russo, M. et al. Cancer drug-tolerant persister cells: from biological questions to clinical opportunities. Nature Reviews Cancer 24, 694–717 (2024). 10.1038/s41568-024-00737-z

12 Russo, M. et al. Adaptive mutability of colorectal cancers in response to targeted therapies. Science 366, 1473–1480 (2019). 10.1126/science.aav4474

13 Russo, M. et al. A modified fluctuation-test framework characterizes the population dynamics and mutation rate of colorectal cancer persister cells. Nature Genetics 54, 976–984 (2022). 10.1038/s41588-022-01105-z

14 Noronha, A. et al. AXL and error-prone DNA replication confer drug resistance and offer strategies to treat EGFR-mutant lung cancer. Cancer Discov (2022). 10.1158/2159-8290.Cd-22-0111

15 Phan, T. G. & Croucher, P. I. The dormant cancer cell life cycle. Nature Reviews Cancer 20, 398–411 (2020). 10.1038/s41568-020-0263-0

16 Cabanos, H. F. & Hata, A. N. Emerging Insights into Targeted Therapy-Tolerant Persister Cells in Cancer. Cancers (Basel) 13 (2021). 10.3390/cancers13112666

17 Oren, Y. et al. Cycling cancer persister cells arise from lineages with distinct programs. Nature 596, 576–582 (2021). 10.1038/s41586-021-03796-6

18 Pisco, A. O. & Huang, S. Non-genetic cancer cell plasticity and therapy-induced stemness in tumour relapse: ‘What does not kill me strengthens me’. British Journal of Cancer 112, 1725–1732 (2015). 10.1038/bjc.2015.146

19 Pisco, A. O. et al. Non-Darwinian dynamics in therapy-induced cancer drug resistance. Nat Commun 4, 2467 (2013). 10.1038/ncomms3467

20 Ramirez, M. et al. Diverse drug-resistance mechanisms can emerge from drug-tolerant cancer persister cells. Nature Communications 7, 10690 (2016). 10.1038/ncomms10690

21 Shen, S., Vagner, S. & Robert, C. Persistent Cancer Cells: The Deadly Survivors. Cell 183, 860–874 (2020). 10.1016/j.cell.2020.10.027

22 Zhao, B. et al. Exploiting Temporal Collateral Sensitivity in Tumor Clonal Evolution. Cell 165, 234–246 (2016). 10.1016/j.cell.2016.01.045

23 Pu, Y. et al. Drug-tolerant persister cells in cancer: the cutting edges and future directions. Nature Reviews Clinical Oncology (2023). 10.1038/s41571-023-00815-5

24 Rodriguez, R., Schreiber, S. L. & Conrad, M. Persister cancer cells: Iron addiction and vulnerability to ferroptosis. Mol Cell 82, 728–740 (2022). 10.1016/j.molcel.2021.12.001

25 You, J. H., Lee, J. & Roh, J. L. Mitochondrial pyruvate carrier 1 regulates ferroptosis in drug-tolerant persister head and neck cancer cells via epithelial-mesenchymal transition. Cancer Lett 507, 40–54 (2021). 10.1016/j.canlet.2021.03.013

26 Hangauer, M. J. et al. Drug-tolerant persister cancer cells are vulnerable to GPX4 inhibition. Nature 551, 247–250 (2017). 10.1038/nature24297

27 Kalkavan, H. et al. Sublethal cytochrome c release generates drug-tolerant persister cells. Cell 185, 3356–3374.e3322 (2022). 10.1016/j.cell.2022.07.025

28 Viswanathan, V. S. et al. Dependency of a therapy-resistant state of cancer cells on a lipid peroxidase pathway. Nature 547, 453–457 (2017). 10.1038/nature23007

29 Kawakami, R. et al. ALDH1A3-mTOR axis as a therapeutic target for anticancer drug-tolerant persister cells in gastric cancer. Cancer Sci 111, 962–973 (2020). 10.1111/cas.14316

30 Ravindran Menon, D. et al. A stress-induced early innate response causes multidrug tolerance in melanoma. Oncogene 34, 4448–4459 (2015). 10.1038/onc.2014.372

31 Raha, D. et al. The cancer stem cell marker aldehyde dehydrogenase is required to maintain a drug-tolerant tumor cell subpopulation. Cancer Res 74, 3579–3590 (2014). 10.1158/0008-5472.Can-13-3456

32 Vinogradova, M. et al. An inhibitor of KDM5 demethylases reduces survival of drug-tolerant cancer cells. Nat Chem Biol 12, 531–538 (2016). 10.1038/nchembio.2085

33 Dhanyamraju, P. K., Schell, T. D., Amin, S. & Robertson, G. P. Drug-Tolerant Persister Cells in Cancer Therapy Resistance. Cancer Res 82, 2503–2514 (2022). 10.1158/0008-5472.Can-21-3844

34 Guler, G. D. et al. Repression of Stress-Induced LINE-1 Expression Protects Cancer Cell Subpopulations from Lethal Drug Exposure. Cancer Cell 32, 221–237.e213 (2017). 10.1016/j.ccell.2017.07.002

35 Aissa, A. F. et al. Single-cell transcriptional changes associated with drug tolerance and response to combination therapies in cancer. Nat Commun 12, 1628 (2021). 10.1038/s41467-021-21884-z

36 Davis, W. J. H., Drummond, C. J., Diermeier, S. & Reid, G. The Potential Links between lncRNAs and Drug Tolerance in Lung Adenocarcinoma. Genes 15, 906 (2024).

37 Mattick, J. S. et al. Long non-coding RNAs: definitions, functions, challenges and recommendations. Nature Reviews Molecular Cell Biology (2023). 10.1038/s41580-022-00566-8

38 Statello, L., Guo, C. J., Chen, L. L. & Huarte, M. Gene regulation by long non-coding RNAs and its biological functions. Nat Rev Mol Cell Biol 22, 96–118 (2021). 10.1038/s41580-020-00315-9

39 Huarte, M. The emerging role of lncRNAs in cancer. Nature Medicine 21, 1253–1261 (2015). 10.1038/nm.3981

40 Yu, J. et al. Induced pluripotent stem cell lines derived from human somatic cells. Science 318, 1917–1920 (2007). 10.1126/science.1151526

41 Searcy, M. B. et al. PAX3-FOXO1 dictates myogenic reprogramming and rhabdomyosarcoma identity in endothelial progenitors. Nature Communications 14, 7291 (2023). 10.1038/s41467-023-43044-1

42 Dull, T. et al. A third-generation lentivirus vector with a conditional packaging system. J Virol 72, 8463–8471 (1998). 10.1128/jvi.72.11.8463-8471.1998

43 Tripathi, V. et al. Long noncoding RNA MALAT1 controls cell cycle progression by regulating the expression of oncogenic transcription factor B-MYB. PLoS Genet 9, e1003368 (2013). 10.1371/journal.pgen.1003368

44 Barry, G. et al. The long non-coding RNA NEAT1 is responsive to neuronal activity and is associated with hyperexcitability states. Scientific Reports 7, 40127 (2017). 10.1038/srep40127

45 Livak, K. J. & Schmittgen, T. D. Analysis of relative gene expression data using real-time quantitative PCR and the 2(-Delta Delta C(T)) Method. Methods 25, 402–408 (2001). 10.1006/meth.2001.1262

46 Shin, M., Meda Krishnamurthy, P., Devi, G. & Watts, J. K. Quantification of Antisense Oligonucleotides by Splint Ligation and Quantitative Polymerase Chain Reaction. Nucleic Acid Ther 32, 66–73 (2022). 10.1089/nat.2021.0040

47 Pierce, A. J., Johnson, R. D., Thompson, L. H. & Jasin, M. XRCC3 promotes homology-directed repair of DNA damage in mammalian cells. Genes & development 13, 2633–2638 (1999). 10.1101/gad.13.20.2633

48 Richardson, C., Moynahan, M. E. & Jasin, M. Double-strand break repair by interchromosomal recombination: suppression of chromosomal translocations. Genes Dev 12, 3831–3842 (1998). 10.1101/gad.12.24.3831

49 Chatterjee, A., Ahn, A., Rodger, E. J., Stockwell, P. A. & Eccles, M. R. in Gene Expression Analysis: Methods and Protocols (eds Nalini Raghavachari & Natàlia Garcia-Reyero) 35–80 (Springer New York, 2018).

50 Gimenez, G., Stockwell, P. A., Rodger, E. J. & Chatterjee, A. in Oral Biology: Molecular Techniques and Applications (eds Gregory J. Seymour, Mary P. Cullinan, Nicholas C. K. Heng, & Paul R. Cooper) 249–278 (Springer US, 2023).

51 Dobin, A. et al. STAR: ultrafast universal RNA-seq aligner. Bioinformatics 29, 15–21 (2013). 10.1093/bioinformatics/bts635

52 Liao, Y., Smyth, G. K. & Shi, W. featureCounts: an efficient general purpose program for assigning sequence reads to genomic features. Bioinformatics 30, 923–930 (2013). 10.1093/bioinformatics/btt656

53 Love, M. I., Huber, W. & Anders, S. Moderated estimation of fold change and dispersion for RNA-seq data with DESeq2. Genome Biology 15, 550 (2014). 10.1186/s13059-014-0550-8

54 Yu, G., Wang, L. G., Han, Y. & He, Q. Y. clusterProfiler: an R package for comparing biological themes among gene clusters. Omics 16, 284–287 (2012). 10.1089/omi.2011.0118

55 Salgia, R. & Kulkarni, P. The Genetic/Non-genetic Duality of Drug ‘Resistance’ in Cancer. Trends in Cancer 4, 110–118 (2018). 10.1016/j.trecan.2018.01.001

56 Kumar, D. et al. LncRNA Malat1 suppresses pyroptosis and T cell-mediated killing of incipient metastatic cells. Nature Cancer 5, 262–282 (2024). 10.1038/s43018-023-00695-9

57 West, Jason A. et al. The Long Noncoding RNAs NEAT1 and MALAT1 Bind Active Chromatin Sites. Molecular Cell 55, 791–802 (2014). 10.1016/j.molcel.2014.07.012

58 Reid, G. et al. Potent subunit-specific effects on cell growth and drug sensitivity from optimised siRNA-mediated silencing of ribonucleotide reductase. J RNAi Gene Silencing 5, 321–330 (2009).

59 Shaffer, S. M. et al. Rare cell variability and drug-induced reprogramming as a mode of cancer drug resistance. Nature 546, 431–435 (2017). 10.1038/nature22794

60 Huang, J. et al. Targeting MALAT1 induces DNA damage and sensitize non-small cell lung cancer cells to cisplatin by repressing BRCA1. Cancer Chemother Pharmacol 86, 663–672 (2020). 10.1007/s00280-020-04152-7

61 Yadav, A. et al. Targeting MALAT1 Augments Sensitivity to PARP Inhibition by Impairing Homologous Recombination in Prostate Cancer. Cancer Research Communications 3, 2044–2061 (2023). 10.1158/2767-9764.Crc-23-0089

62 Hu, Y. et al. Targeting the MALAT1/PARP1/LIG3 complex induces DNA damage and apoptosis in multiple myeloma. Leukemia 32, 2250–2262 (2018). 10.1038/s41375-018-0104-2

63 Soria, J.-C. et al. Osimertinib in Untreated EGFR-Mutated Advanced Non–Small-Cell Lung Cancer. New England Journal of Medicine 378, 113–125 (2017). 10.1056/NEJMoa1713137

64 Jänne, P. A. et al. Adagrasib in Non Small-Cell Lung Cancer Harboring a *KRAS*G12C Mutation. New England Journal of Medicine 387, 120–131 (2022). 10.1056/NEJMoa2204619

65 Hong, D. S. et al. KRAS(G12C) Inhibition with Sotorasib in Advanced Solid Tumors. N Engl J Med 383, 1207–1217 (2020). 10.1056/NEJMoa1917239

66 Planchard, D. et al. Osimertinib with or without Chemotherapy in EGFR-Mutated Advanced NSCLC. N Engl J Med 389, 1935–1948 (2023). 10.1056/NEJMoa2306434

67 Mok, T. S. K. et al. KRYSTAL-12: Phase 3 study of adagrasib versus docetaxel in patients with previously treated advanced/metastatic non-small cell lung cancer (NSCLC) harboring a KRASG12C mutation. Journal of Clinical Oncology 42, LBA8509–LBA8509 (2024). 10.1200/JCO.2024.42.17_suppl.LBA8509

68 Mok, T. S. et al. Osimertinib or Platinum Pemetrexed in *EGFR* T790M Positive Lung Cancer. New England Journal of Medicine 376, 629–640 (2017). 10.1056/NEJMoa1612674

69 Yuan, C. et al. DNA damage/cGAS-triggered up-regulation of MALAT1 promotes undesirable inflammatory responses in radiotherapy of cancer. Biochem Biophys Res Commun 528, 746–752 (2020). 10.1016/j.bbrc.2020.05.064

70 Wang, Y.-L. et al. DNA damage-induced paraspeckle formation enhances DNA repair and tumor radioresistance by recruiting ribosomal protein P0. Cell Death & Disease 13, 709 (2022). 10.1038/s41419-022-05092-1

71 Blume, C. J. et al. p53-dependent non-coding RNA networks in chronic lymphocytic leukemia. Leukemia 29, 2015–2023 (2015). 10.1038/leu.2015.119

72 Isozaki, H. et al. Therapy-induced APOBEC3A drives evolution of persistent cancer cells. Nature (2023). 10.1038/s41586-023-06303-1

73 Luo, J. et al. LncRNA MALAT-1 modulates EGFR-TKI resistance in lung adenocarcinoma cells by downregulating miR-125. Discov Oncol 15, 379 (2024). 10.1007/s12672-024-01133-7

74 Wang, S., Hu, C., Wu, F. & He, S. Rab25 GTPase: Functional roles in cancer. Oncotarget 8, 64591–64599 (2017). 10.18632/oncotarget.19571

75 Ji, P. et al. MALAT-1, a novel noncoding RNA, and thymosin beta4 predict metastasis and survival in early-stage non-small cell lung cancer. Oncogene 22, 8031–8041 (2003). 10.1038/sj.onc.1206928

76 Arun, G., Aggarwal, D. & Spector, D. L. MALAT1 Long Non-Coding RNA: Functional Implications. Non-Coding RNA 6 (2020).

77 Amodio, N. et al. MALAT1: a druggable long non-coding RNA for targeted anti-cancer approaches. Journal of Hematology & Oncology 11, 63 (2018). 10.1186/s13045-018-0606-4

78 Schmidt, L. H. et al. The long noncoding MALAT-1 RNA indicates a poor prognosis in non-small cell lung cancer and induces migration and tumor growth. J Thorac Oncol 6, 1984–1992 (2011). 10.1097/JTO.0b013e3182307eac

79 Gutschner, T. et al. The Noncoding RNA MALAT1 Is a Critical Regulator of the Metastasis Phenotype of Lung Cancer Cells. Cancer Research 73, 1180–1189 (2013). 10.1158/0008-5472.Can-12-2850

80 Fang, Z., Chen, W., Yuan, Z., Liu, X. & Jiang, H. LncRNA-MALAT1 contributes to the cisplatin-resistance of lung cancer by upregulating MRP1 and MDR1 via STAT3 activation. Biomed Pharmacother 101, 536–542 (2018). 10.1016/j.biopha.2018.02.130

81 Temprine, K. et al. Regulation of the error-prone DNA polymerase Polκ by oncogenic signaling and its contribution to drug resistance. Sci Signal 13 (2020). 10.1126/scisignal.aau1453

82 Arun, G. et al. Differentiation of mammary tumors and reduction in metastasis upon Malat1 lncRNA loss. Genes Dev 30, 34–51 (2016). 10.1101/gad.270959.115

83 Kim, J. et al. Long noncoding RNA MALAT1 suppresses breast cancer metastasis. Nat Genet 50, 1705–1715 (2018). 10.1038/s41588-018-0252-3

84 Kurppa, K. J. et al. Treatment-Induced Tumor Dormancy through YAP-Mediated Transcriptional Reprogramming of the Apoptotic Pathway. Cancer cell 37, 104–122.e112 (2020). 10.1016/j.ccell.2019.12.006

85 Oh, Y.-T., Chen, Z., Wang, D., Ramalingam, S. S. & Sun, S.-Y. Induction of IL6/STAT3-dependent TRAIL expression that contributes to the therapeutic efficacy of osimertinib in EGFR mutant NSCLC cells. Oncogene 44, 2315–2327 (2025). 10.1038/s41388-025-03381-5

86 Arasada, R. R. et al. Notch3-dependent β-catenin signaling mediates EGFR TKI drug persistence in EGFR mutant NSCLC. Nature Communications 9, 3198 (2018). 10.1038/s41467-018-05626-2

87 Kim, S. M. et al. Activation of IL-6R/JAK1/STAT3 Signaling Induces De Novo Resistance to Irreversible EGFR Inhibitors in Non–Small Cell Lung Cancer with T790M Resistance Mutation. Molecular Cancer Therapeutics 11, 2254–2264 (2012). 10.1158/1535-7163.Mct-12-0311

88 Johnson, D. E., O’Keefe, R. A. & Grandis, J. R. Targeting the IL-6/JAK/STAT3 signalling axis in cancer. Nature Reviews Clinical Oncology 15, 234–248 (2018). 10.1038/nrclinonc.2018.8

89 Kubala, M. H. & DeClerck, Y. A. The plasminogen activator inhibitor-1 paradox in cancer: a mechanistic understanding. Cancer Metastasis Rev 38, 483–492 (2019). 10.1007/s10555-019-09806-4

90 Lee, H.-J. et al. Drug Resistance via Feedback Activation of Stat3 in Oncogene-Addicted Cancer Cells. Cancer Cell 26, 207–221 (2014). 10.1016/j.ccr.2014.05.019

91 Zhao, G., Su, Z., Song, D., Mao, Y. & Mao, X. The long noncoding RNA MALAT1 regulates the lipopolysaccharide-induced inflammatory response through its interaction with NF-κB. FEBS Letters 590, 2884–2895 (2016). 10.1002/1873-3468.12315

92 Puthanveetil, P., Chen, S., Feng, B., Gautam, A. & Chakrabarti, S. Long non-coding RNA MALAT1 regulates hyperglycaemia induced inflammatory process in the endothelial cells. J Cell Mol Med 19, 1418–1425 (2015). 10.1111/jcmm.12576

93 Wang, Z. et al. LncRNA MALAT1 promotes gastric cancer progression via inhibiting autophagic flux and inducing fibroblast activation. Cell Death & Disease 12, 368 (2021). 10.1038/s41419-021-03645-4

94 Jadaliha, M. et al. Functional and prognostic significance of long non-coding RNA MALAT1 as a metastasis driver in ER negative lymph node negative breast cancer. Oncotarget 7, 40418–40436 (2016). 10.18632/oncotarget.9622

95 Sillen, M. & Declerck, P. J. A Narrative Review on Plasminogen Activator Inhibitor-1 and Its (Patho)Physiological Role: To Target or Not to Target? International Journal of Molecular Sciences 22, 2721 (2021).

96 Kuilman, T. et al. Oncogene-Induced Senescence Relayed by an Interleukin-Dependent Inflammatory Network. Cell 133, 1019–1031 (2008). 10.1016/j.cell.2008.03.039

97 Coppé, J. P. et al. Senescence-associated secretory phenotypes reveal cell-nonautonomous functions of oncogenic RAS and the p53 tumor suppressor. PLoS Biol 6, 2853–2868 (2008). 10.1371/journal.pbio.0060301

98 Tian, X. et al. β-Actin regulates interleukin 6-induced p21 transcription by interacting with the Rpb5 and Rpb7 subunits of RNA polymerase II. Animal Cells and Systems 20, 282–288 (2016). 10.1080/19768354.2016.1224204

99 Moran, D. M., Mattocks, M. A., Cahill, P. A., Koniaris, L. G. & McKillop, I. H. Interleukin-6 mediates G(0)/G(1) growth arrest in hepatocellular carcinoma through a STAT 3-dependent pathway. J Surg Res 147, 23–33 (2008). 10.1016/j.jss.2007.04.022

100 Flørenes, V. A. et al. Interleukin-6 dependent induction of the cyclin dependent kinase inhibitor p21WAF1/CIP1 is lost during progression of human malignant melanoma. Oncogene 18, 1023–1032 (1999). 10.1038/sj.onc.1202382

101 Eißmann, M. et al. Loss of the abundant nuclear non-coding RNA MALAT1 is compatible with life and development. RNA Biol 9, 1076–1087 (2012). 10.4161/rna.21089

102 Zhang, B. et al. The lncRNA Malat1 is dispensable for mouse development but its transcription plays a cis-regulatory role in the adult. Cell Rep 2, 111–123 (2012). 10.1016/j.celrep.2012.06.003

103 Crooke, S. T., Baker, B. F., Crooke, R. M. & Liang, X.-h. Antisense technology: an overview and prospectus. Nature Reviews Drug Discovery 20, 427–453 (2021). 10.1038/s41573-021-00162-z

104 Roberts, T. C., Langer, R. & Wood, M. J. A. Advances in oligonucleotide drug delivery. Nature Reviews Drug Discovery 19, 673–694 (2020). 10.1038/s41573-020-0075-7

105 Kulkarni, J. A. et al. The current landscape of nucleic acid therapeutics. Nature Nanotechnology 16, 630–643 (2021). 10.1038/s41565-021-00898-0

106 Wang, X. et al. Preparation of selective organ-targeting (SORT) lipid nanoparticles (LNPs) using multiple technical methods for tissue-specific mRNA delivery. Nature Protocols 18, 265–291 (2023). 10.1038/s41596-022-00755-x

107 Cheng, Q. et al. Selective organ targeting (SORT) nanoparticles for tissue-specific mRNA delivery and CRISPR–Cas gene editing. Nature Nanotechnology 15, 313–320 (2020). 10.1038/s41565-020-0669-6

108 Lauffer, M. C., van Roon-Mom, W. & Aartsma-Rus, A. Possibilities and limitations of antisense oligonucleotide therapies for the treatment of monogenic disorders. Communications Medicine 4, 6 (2024). 10.1038/s43856-023-00419-1

109 Wiesenfarth, M. et al. Effects of tofersen treatment in patients with SOD1-ALS in a “real-world” setting - a 12-month multicenter cohort study from the German early access program. eClinicalMedicine 69 (2024). 10.1016/j.eclinm.2024.102495

110 Brown, J. A., Valenstein, M. L., Yario, T. A., Tycowski, K. T. & Steitz, J. A. Formation of triple-helical structures by the 3’-end sequences of MALAT1 and MENβ noncoding RNAs. Proc Natl Acad Sci U S A 109, 19202–19207 (2012). 10.1073/pnas.1217338109

111 Abulwerdi, F. A. et al. Selective Small-Molecule Targeting of a Triple Helix Encoded by the Long Noncoding RNA, MALAT1. ACS Chemical Biology 14, 223–235 (2019). 10.1021/acschembio.8b00807

112 Childs-Disney, J. L. et al. Targeting RNA structures with small molecules. Nature Reviews Drug Discovery 21, 736–762 (2022). 10.1038/s41573-022-00521-4

113 Donlic, A. et al. Discovery of Small Molecule Ligands for MALAT1 by Tuning an RNA-Binding Scaffold. Angew Chem Int Ed Engl 57, 13242–13247 (2018). 10.1002/anie.201808823

114 Chen, W., Mook, R. A., Jr., Premont, R. T. & Wang, J. Niclosamide: Beyond an antihelminthic drug. Cell Signal 41, 89–96 (2018). 10.1016/j.cellsig.2017.04.001

115 Burock, S. et al. Phase II trial to investigate the safety and efficacy of orally applied niclosamide in patients with metachronous or sychronous metastases of a colorectal cancer progressing after therapy: the NIKOLO trial. BMC Cancer 18, 297 (2018). 10.1186/s12885-018-4197-9

116 Zablowsky, N. et al. High Throughput FISH Screening Identifies Small Molecules That Modulate Oncogenic lncRNA MALAT1 via GSK3B and hnRNPs. Noncoding RNA 9 (2023). 10.3390/ncrna9010002

