## Supplemental information for "Elevated Expression of *MALAT1* Contributes to the Survival of Drug-Tolerant Persister Cells Following Targeted Therapy in Lung Adenocarcinoma"

**Supplementary Figures:**

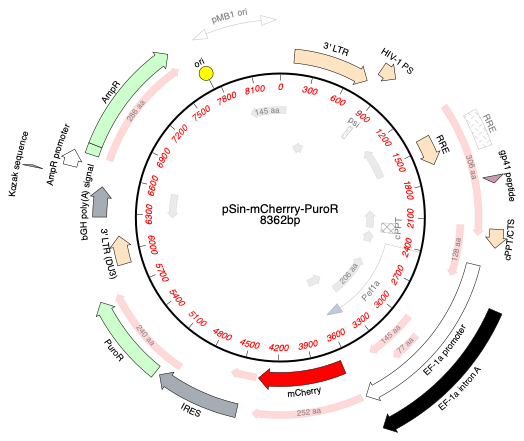

**Supplementary Figure 1:**  Plasmid Map for pSin-mCherry-PuroR Plasmid. Map generated in MacVector (version 18.6.0).

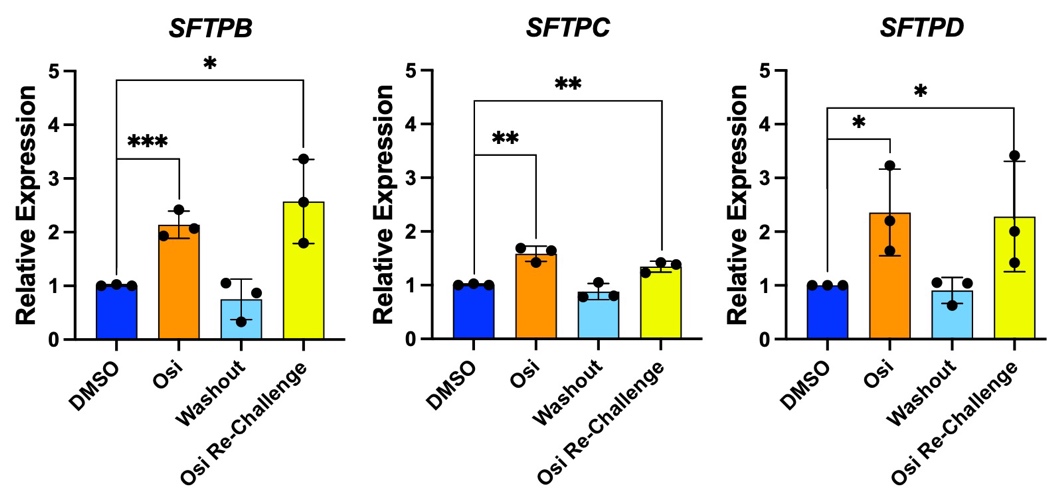

**Supplementary Figure 2:** Relative expression of *SFTPB*, *SFTPC* and *SFTPD* in osimertinib treatment, washout and re-challenge experiments relative to solvent controls in PC9 cells by RT-qPCR. control. Unpaired, one-tailed t-tests were performed to test for significance. *=p ≤ 0.05, **=p ≤ 0.01, ***=p ≤ 0.00. Data is mean ± standard deviation (n=3).

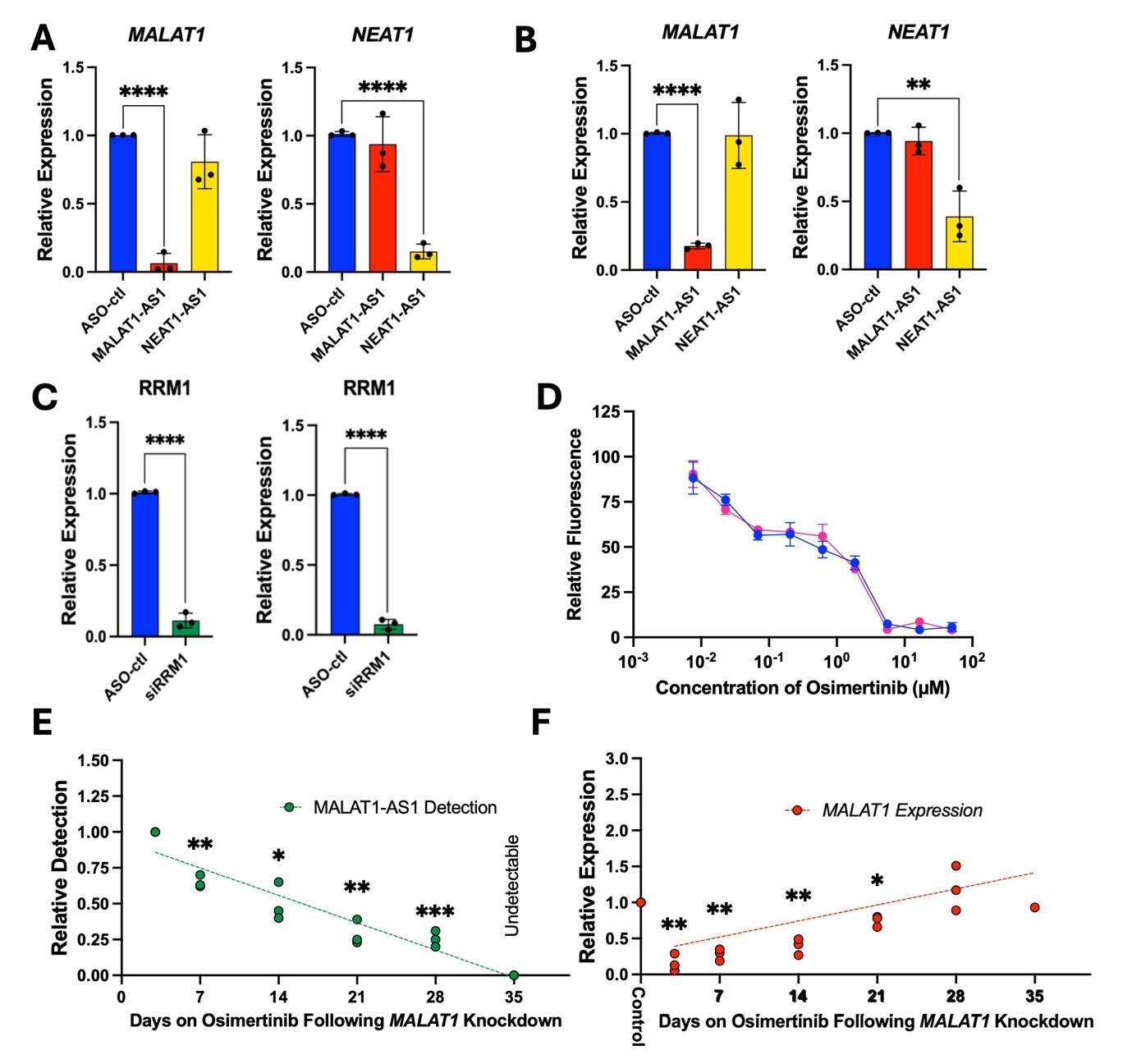

**Supplementary Figure 3:** PC9 cells **(A)**, or H358 cells **(B)** were reverse transfected for 48 hours with a control ASO (ASO-control), MALAT1-AS1 or NEAT1-AS1. RT-qPCR was then performed using primers for *MALAT1* (left) or *NEAT1* (right). **(C)** RT-qPCR for *RRM1* positive transfection control for knockdown experiments in PC9 cells (left) and H358 cells (right). **(D)** Proliferation assay comparing viability of WT-PC9 cells with lentivirally transduced pSin-mCherry-PuroR expressing PC9 cells. Error bars represent the mean ± standard deviation of 3 internal technical replicates. Representative figure shown (n=3). **(E and F)** PC9 cells were reverse transfected with MALAT1-AS1 and treated with 2 µM osimertinib for five weeks. SplintR hybridisation, ligation and TaqMan qPCR were performed **(E)** and results were normalised to the highest ASO-concentration (Day 3). **(F)** SYBR-based RT-qPCR detection of *MALAT1*. SYBR-based RT-qPCR was normalised to UBC as a housekeeping control, and to the DMSO/ASO-control. Linear regressions (dashed lines) were performed in GraphPad PRISM. Unpaired, one-tailed t-tests were performed to test for significance. *=p ≤ 0.05, **=p ≤ 0.01, ***=p ≤ 0.001, ****=p ≤ 0.0001. Data is mean ± standard deviation (n=3).

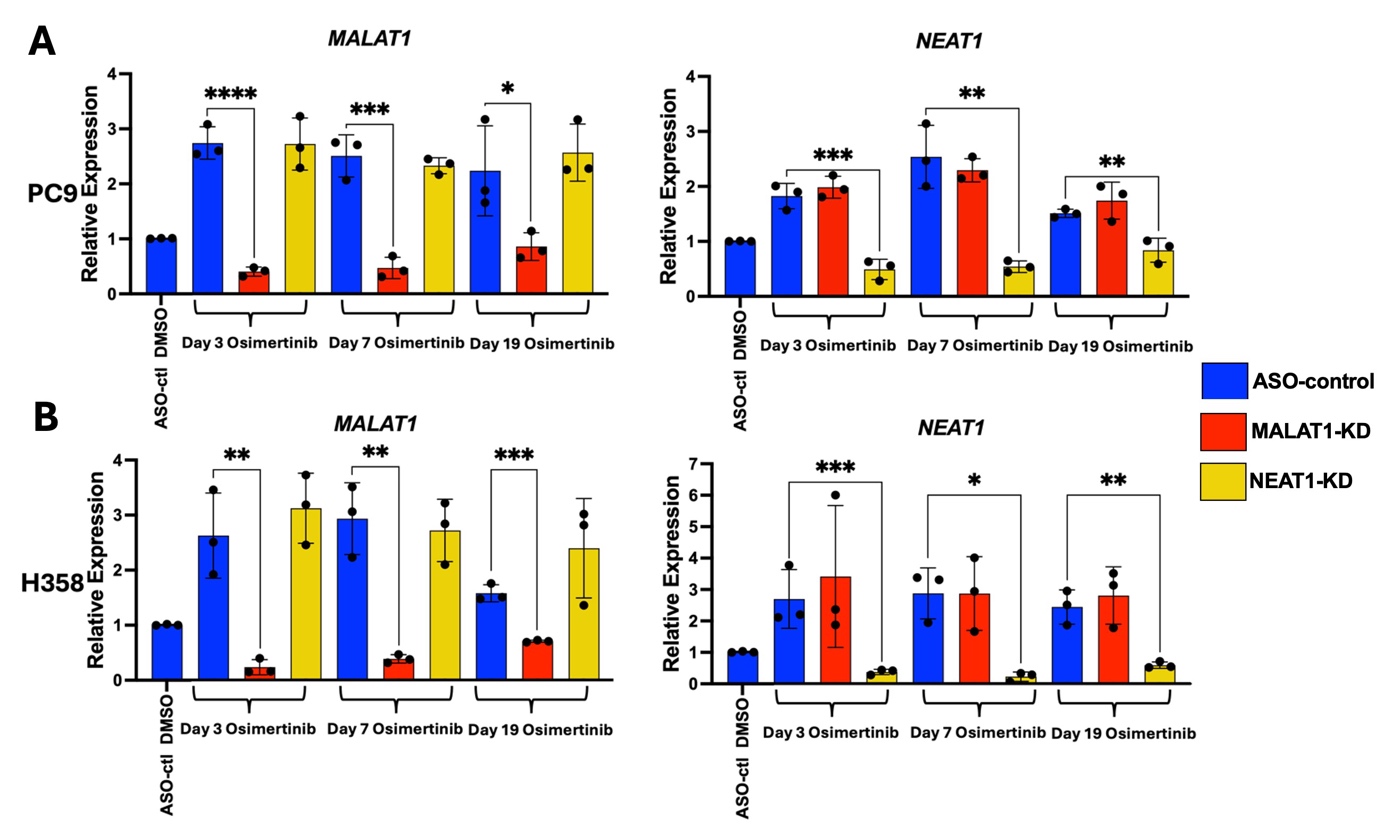

**Supplementary Figure 4:** ASO mediated knockdown of *MALAT1* and *NEAT1* in DTPs treated for 3, 7, or 19 days in PC9 cells **(A)** or H358 cells **(B)**. Relative expression of *MALAT1* or *NEAT1* in relative to solvent controls targeted therapy treated PC9 or H358 cells was determined by RT-qPCR. Unpaired, one-tailed t-tests were performed to test for significance. *=p ≤ 0.05, **=p ≤ 0.01, ***=p ≤ 0.001, ****=p ≤ 0.0001. Data is mean ± standard deviation (n=3).

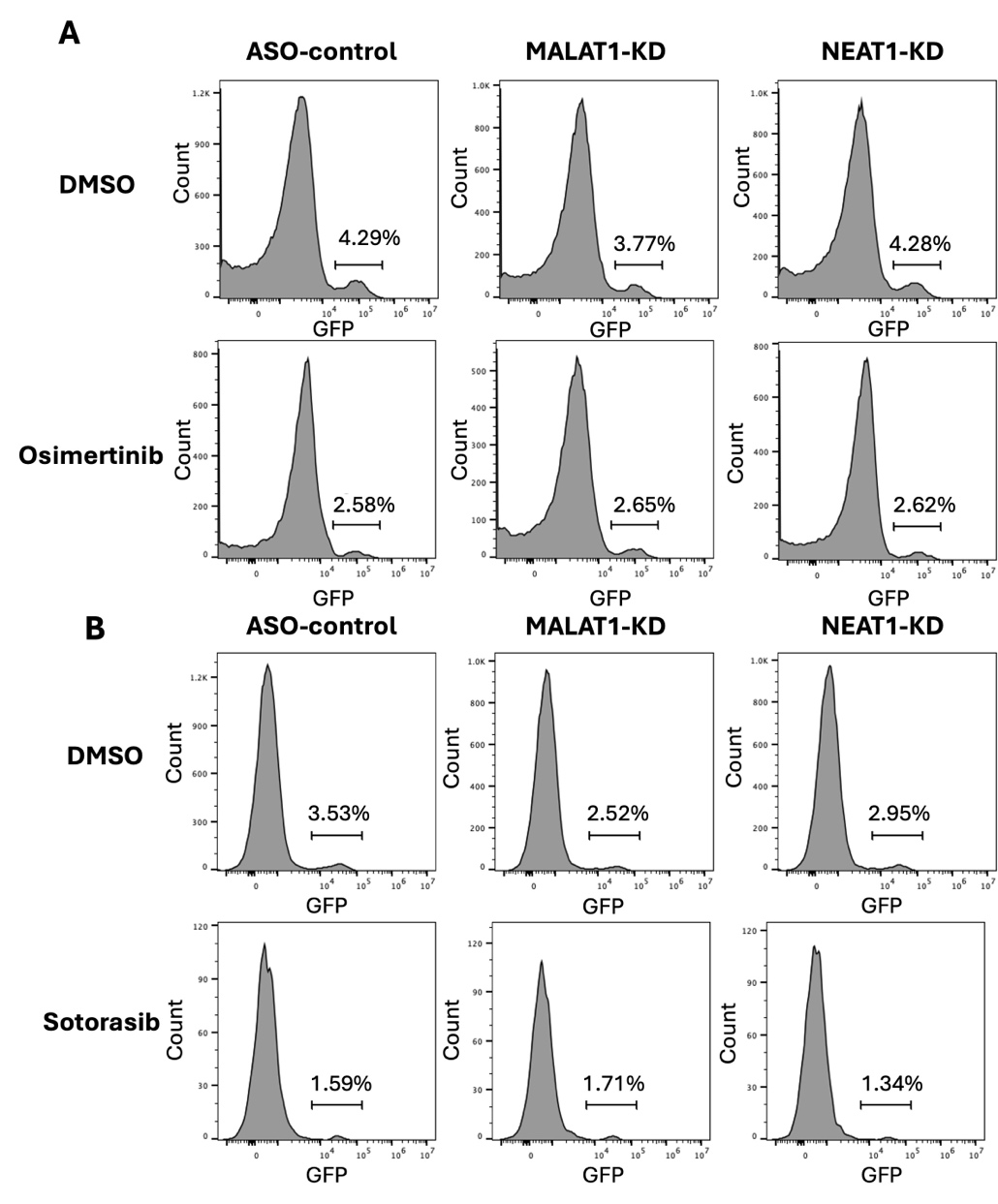

**Supplementary Figure 5:** Representative Flow Cytometry Plots for the Homologous Recombination Assay in PC9 and H358 Cells. PC9 cells **(A)** or H358 cells **(B)** were reverse transfected with an ASO-control, *MALAT1* or *NEAT1* ASOs and treated with a solvent control or targeted therapy as indicated for 72-hours. Cells positive for homologous recombination were first obtained by gating the live population of cells (Zombie-NIR) and then by cells co-transfected with a constitutive mCherry plasmid to determine transfection efficiency. Finally, the percent of live, positively transfected cells able to recombine the pDRGFP plasmid via homologous recombination was quantified in the GFP channel and is shown above. Live/dead, mCherry and GFP gates were determined via positive and negative single colour controls. All gates remained consistent between conditions within each experiment. Plots were generated via FlowJo (v10.10).

**Supplementary Tables:**

Supplementary Table 1: Sequencing Primers Used for Cloning pSin-mCherry-PuroR Plasmid

| Primer | Sequence (5’- 3’) |
| --- | --- |
| EF1A-Fwd | TCAAGCCTCAGACAGTGGTTC |
| IRES-Rev | GCATTCCTTTGGCGAGAG |

Supplementary Table 2: ASO and siRNA Sequences and Modifications

| ASO | Sequence | Sequence with Modifications (5’ - 3’) |
| --- | --- | --- |
| ASO-control | CCTTCCTGAAGGTTCTCC | moC*moC*moT*moT*moC*moC*moT*moG*moA*moA*moG*moT*moT*moC*moT*moC*moC* |
| NEAT1-AS1 | ATCACACATGTAGTAAAGGC | moA*moT*moC*moA*moC*dA*dC*dA*dT*dG*d  T*dA*dG*dT*dA*moA*moA*moG*moG*moC |
| NEAT1-AS2 | TCGCTCATGATTTTCAATCA | moT*moC*moG*moC*moT*dC*dA*dT*dG*dA*d  T*dT*dT*dT*dC*moA*moA*moT*moC*moA |
| NEAT1-AS3 | ATCACACATGTAGTAAAGGC | moA*moT*moC*moA*moC*dA*dC*dA*dT*dG*d  T*dA*dG*dT*dA*moA*moA*moG*moG*moC |
| NEAT1-AS4 | ATCATCCCCAAGTCATTGGT | moA*moT*moC*moA*moT*dC*dC*dC*dC*dA*d  A*dG*dT*dC*dA*moT*moT*moG*moG*moT |
| MALAT1-AS1 | GGGAGTTACTTGCCAACTTG | moG*moG*moG*moA*moG*dT*dT*dA*dC*dT*d  T*dG*dC*dC*dA*moA*moC*moT*moT*moG |
| MALAT1-AS2 | ATGGAGGTATGACATATAAT | moA*moT*moG*moG*moA*dG*dG*dT*dA*dT*d  G*dA*dC*dA*dT*moA*moT*moA*moA*moT |
| siRRM1 | GCAAACUCACUAGUAUGCACUUCUA | mGmCmAmAmAmCUCACUAGU AUGCAmCmUmUmCmUmA |

ASOs were modified with 2’-O-Methyl RNA, (mN), 2’methoxyethyl RNA (moN) or DNA (dN) nucleotides and phosphorothioate linkages (*) as indicated.

Supplementary Table 3: Primers Used for RT-qPCR

| Gene | Forward Primer (5’ - 3’) | Reverse Primer (5’ - 3’) | Reference |
| --- | --- | --- | --- |
| *ANRIL* | GACCAGTGACTTAGATTGGCTTC | TCAAGTGTGTAGAGCTTCGTATCC | ^117^ |
| *AQP4* | TCAGCATCGCCAAGTCTGTC | CTGGGAGGTGTGACCAGATAG |  |
| *BRCA1* | AGAAACCACCAAGGTCCAAAG | GGGCCCATAGCAACAGATTT | ^14^ |
| *BRCA2* | AGGACTTGCCCCTTTCGTCTA | TGCAGCAATTAACATATGAGG | ^14^ |
| *EXO1* | CCTCGTGGCTCCCTATGAAG | AGGAGATCCGAGTCCTCTGTAA | ^14^ |
| *IL6* | GGATTCAATGAGGAGACTTGCC | TGGCATTTGTGGTTGGGTCA | ^118^ |
| *MALAT1* | CGCATTTACTAAACGCAGAC | TCTCTATTCTTTTCTTCGCC |  |
| *MLH1* | CAGAGCTTGGAGGGGGATA | TTTCGGGAATCATCTTCCAC | ^14^ |
| *MSH6* | GGGGCAAGTCTACGCTTATG | CACACTTCAGCAGGGACGTA | ^14^ |
| *NEAT1* | GCTGGACCTTTCATGTAACGGG | TGAACTCTGCCGGTACAGGGAA |  |
| *PMS2* | TTTGCCGACCTAACTCAGGTT | CGATGCGTGGCAGGTAGAA | ^14^ |
| *POLA1* | ACGCCAGGATGATGACTGGA | GTCACTGCGAGCTTCTTTACAT | ^14^ |
| *POLD1* | CAGTGCCAAGGTGGTGTATGG | CTTGCTGATAAGCAGGTATGGG | ^14^ |
| *POLE* | TTGCGACCAGAAAGGGTTGT | TGATTTGGCAAGTCCAGATCCT | ^14^ |
| *POLι* | ACAAACCGGGATTTCCTACC | TCACACTTCCTTTCCCTTGAA | ^14^ |
| *POLκ* | TGAGGGACAATCCAGAATTGAAG | CTGCACGAACACCAAATCTCC | ^14^ |
| *POLλ* | GATGAAGGCATGGACTATGAGC | TGAATCCAGCTACATCCACCAG | ^14^ |
| *REV1* | ACCGAAGAGGAGCACAAAGA | CCATTCCATTTCCCTGAAGA | ^14^ |
| *RRM1* | GGCAAACTCACTAGTATGCACTTC | AAATAATACATCCCAGTCTTCAAACC |  |
| *SERPINE1* | TGCCCATGATGGCTCAGA | GCAGTTCCAGGATGTCGTAGTAATG | ^119^ |
| *SFTPB* | GCAACGTCCTCCCCTTGAAG | AGTCAGTCTGGTTCTGGAAGTAG | ^7^ |
| *SFTPC* | CACCTGAAACGCCTTCTTATCG | TTTCTGGCTCATGTGGAGACC | ^7^ |
| *SFTPD* | AAGCAGGGGAACATAGGACCT | ACACCTCGCTCTCCCTTAGG | ^7^ |
| *UBC* | GCAAAGATCCAAGATAAGGAA | GGACCAAGTGCAGAGTGGAC |  |
| *UFC1* | TCCAACCTGAGTGACATAGCGA | CTGACCTCCAACTCCAACGAAT | ^120^ |

Supplementary Table 4: Sequence Information for SplintR Assays

| Oligonucleotide | Sequence (5’- 3’) |
| --- | --- |
| Probe A | CTCGACCTCTCTATGGGCAGTCACGACAGCAAGTTGGCA |
| Probe B | pAGTAACTCCCCTGAGTCGGAGACACGCAGGGCTTAA |
| Forward Primer | GCTCGACCTCTCTATGGGC |
| Reverse Primer | TTAAGCCCTGCGTGTCTCC |
| Double-Quenched FAM Probe | /FAM/CTAGCGCGC/ZEN/GACTCCGTCGTG/IABkFQ/ |

Supplementary Table 5: Expression of Differentially Expressed Genes Overlapping at 3, 7, and 19 days of Treatment Following *MALAT1*-KD in DTPs:

| **Gene Name** | **Day 3 Log2FC** | **Day 3 Adj.P-Val.** | **Day 7 Log2FC** | **Day 7 Adj.P-Val.** | **Day 19 Log2FC** | **Day 19 Adj.P-Val.** | **Av. Log2FC** |
| --- | --- | --- | --- | --- | --- | --- | --- |
| ACP5 | 3.375292975 | 0.001864943 | 3.973802527 | 0.000262232 | 3.374277655 | 0.002229045 | 3.574457719 |
| ACTBL2 | 1.72710544 | 0.002907499 | 1.715717003 | 0.006368045 | 2.094985193 | 0.000266213 | 1.845935879 |
| ADAM32 | 2.185749631 | 0.000366422 | 1.828075678 | 0.007229753 | 1.8422747 | 0.006415028 | 1.952033336 |
| ADAM8 | 1.479513227 | 0.000166136 | 1.331754369 | 0.001765313 | 1.294006489 | 0.001999824 | 1.368424695 |
| ADAMTS14 | 1.751833458 | 0.034375625 | 1.762151922 | 0.047755695 | 2.332629802 | 0.003060302 | 1.948871727 |
| ADAMTS3 | -2.026024008 | 1.26344E-07 | -2.115342119 | 2.22835E-08 | -2.119502738 | 1.43588E-08 | -2.08695629 |
| ADAMTS6 | 1.924561964 | 3.05445E-07 | 1.744908421 | 1.03911E-05 | 2.095759774 | 3.03685E-08 | 1.921743387 |
| ALOX5 | 1.670225472 | 0.000854138 | 1.423029107 | 0.013795545 | 2.517582508 | 1.07348E-06 | 1.870279029 |
| ALPG | -2.808119475 | 0.000813479 | -2.25145491 | 0.014812619 | -2.11068158 | 0.020530336 | -2.39008532 |
| ALPK2 | -1.87108848 | 0.048046058 | -1.966879153 | 0.033696505 | -3.135024483 | 9.32487E-05 | -2.32433071 |
| ALPP | -2.755925727 | 1.12603E-05 | -1.686514572 | 0.02540413 | -1.84844605 | 0.008952672 | -2.09696212 |
| ANKRD6 | -1.074633642 | 0.00345247 | -1.288236997 | 0.00038218 | -1.862067916 | 1.0399E-08 | -1.40831285 |
| ANKRD65 | 2.719738667 | 9.85731E-06 | 3.025537191 | 2.15147E-06 | 2.757498909 | 2.47576E-05 | 2.834258256 |
| AOX1 | 1.602859473 | 1.4135E-07 | 2.113915402 | 5.77282E-13 | 1.710471566 | 1.2272E-08 | 1.809082147 |
| ARMCX2 | 1.469641044 | 0.002047947 | 1.395043154 | 0.005456223 | 1.557458657 | 0.000832027 | 1.474047618 |
| ARTN | 1.551985087 | 0.007174076 | 1.77000149 | 0.002654119 | 2.441495838 | 6.92817E-06 | 1.921160805 |
| ATG9B | 1.808884248 | 0.014300463 | 2.207723591 | 0.003157983 | 2.486912099 | 0.000339948 | 2.167839979 |
| ATP9A | 1.370768598 | 2.53877E-05 | 1.035008368 | 0.005199229 | 1.187424609 | 0.000572544 | 1.197733858 |
| AVIL | 4.644733321 | 7.68576E-07 | 3.424450139 | 0.001132405 | 4.035222145 | 4.55233E-05 | 4.034801868 |
| BAMBI | -1.292099581 | 0.000387815 | -1.019882968 | 0.012313727 | -1.464885396 | 4.17376E-05 | -1.25895598 |
| BMP6 | -2.02238006 | 0.000477793 | -1.726611367 | 0.00566636 | -2.266923571 | 4.54567E-05 | -2.005305 |
| BSPRY | 1.802187776 | 0.000157076 | 1.598641656 | 0.001981671 | 1.364629728 | 0.009499586 | 1.588486386 |
| C15orf48 | 2.509066894 | 1.13914E-05 | 1.615542691 | 0.017498099 | 2.329417433 | 9.32487E-05 | 2.15134234 |
| C2orf72 | -2.160917946 | 1.94072E-05 | -2.100674951 | 2.27342E-05 | -1.980768176 | 3.29428E-05 | -2.08078702 |
| C6orf15 | 2.879586935 | 0.000598814 | 2.231565296 | 0.027977828 | 2.906804885 | 0.001799587 | 2.672652372 |
| CARD9 | 1.415089324 | 0.001354858 | 1.557202718 | 0.0006918 | 1.901514961 | 1.18186E-05 | 1.624602334 |
| CD68 | 1.711470178 | 1.58497E-07 | 1.201513601 | 0.00139321 | 1.552433788 | 6.92817E-06 | 1.488472522 |
| CDH23 | -1.517400593 | 0.036878499 | -1.557231785 | 0.038695591 | -1.882178795 | 0.005815914 | -1.65227039 |
| CEACAM6 | 5.676920969 | 5.20058E-05 | 4.745548709 | 0.002566559 | 4.440868845 | 0.004310227 | 4.954446174 |
| CFAP58 | 1.167538769 | 0.049053128 | 1.448829768 | 0.010532667 | 1.227143367 | 0.02950815 | 1.281170635 |
| CGB8 | 1.884502859 | 0.02450578 | 1.811392228 | 0.045658279 | 1.962336914 | 0.019791281 | 1.886077334 |
| CHEK2P2 | -6.115926895 | 0.00033365 | -4.704633038 | 0.00249461 | -5.379672059 | 0.000416857 | -5.40007733 |
| CHSY3 | -2.381142431 | 2.6782E-05 | -1.783907504 | 0.003805568 | -2.592560915 | 1.93431E-06 | -2.25253695 |
| CLTRN | 1.965359792 | 0.008553014 | 2.079096884 | 0.008812071 | 2.058649967 | 0.009750861 | 2.034368881 |
| COL4A6 | 2.570574121 | 5.29925E-23 | 2.060984409 | 1.02808E-13 | 2.032697044 | 8.6834E-14 | 2.221418525 |
| CORO6 | 1.241089574 | 0.001138294 | 1.48921854 | 9.96773E-05 | 1.471812056 | 9.94257E-05 | 1.400706723 |
| CPA4 | 2.28865085 | 4.16183E-09 | 1.548606299 | 0.000384884 | 1.570078088 | 0.00020412 | 1.802445079 |
| CPE | -2.021250036 | 3.89858E-08 | -1.66752327 | 1.05041E-05 | -2.03002588 | 1.01875E-08 | -1.9062664 |
| CPNE7 | 1.405747513 | 0.020102822 | 1.846953342 | 0.001723902 | 2.160859569 | 0.000111059 | 1.804520141 |
| CPVL | -2.242802773 | 2.96696E-07 | -2.580997165 | 1.01077E-09 | -3.329013874 | 2.95182E-16 | -2.7176046 |
| CR2 | -1.60043328 | 0.002216009 | -1.369062021 | 0.013151844 | -1.750142746 | 0.000441789 | -1.57321268 |
| CRYBG2 | 1.630926479 | 1.70589E-06 | 1.507922467 | 2.92578E-05 | 1.801582992 | 1.0843E-07 | 1.646810646 |
| CUZD1 | 2.115110734 | 0.000366769 | 1.449844054 | 0.035859126 | 1.536108953 | 0.020064129 | 1.70035458 |
| DAPP1 | 1.242945564 | 0.00015557 | 1.003116475 | 0.006284224 | 1.259049488 | 0.000177107 | 1.168370509 |
| DDX3ILA1 | 2.850252412 | 0.016646342 | 2.979394695 | 0.020417963 | 4.072928142 | 0.000948421 | 3.300858416 |
| DEPTOR | -1.922199144 | 0.004243336 | -1.805236111 | 0.007229753 | -2.434587356 | 0.000119569 | -2.05400754 |
| DIRAS2 | -2.942937638 | 0.002227044 | -2.171462447 | 0.026350493 | -2.87151048 | 0.000798417 | -2.66197019 |
| DYNC2H1 | -1.107967905 | 0.008735062 | -1.117218271 | 0.011486468 | -1.104818469 | 0.009723296 | -1.11000155 |
| EFHD1 | -1.679997866 | 1.08309E-05 | -1.385212016 | 0.000893994 | -1.620508274 | 3.68565E-05 | -1.56190605 |
| EFNB2 | 2.039901912 | 7.22345E-12 | 1.817191682 | 1.54563E-08 | 1.906353254 | 1.91E-09 | 1.921148949 |
| EFNB3 | -1.109480622 | 0.005575779 | -1.099210231 | 0.005947119 | -1.338124692 | 0.000337764 | -1.18227185 |
| ENC1 | 1.170793736 | 1.58497E-07 | 1.292390081 | 7.20194E-09 | 1.283924988 | 8.17027E-09 | 1.249036268 |
| EPDR1 | 4.827779683 | 3.08803E-05 | 4.405858074 | 0.000305765 | 4.881203431 | 2.12032E-05 | 4.704947063 |
| EPHA7 | -3.207615186 | 4.75993E-07 | -1.93999185 | 0.008269175 | -2.194439455 | 0.001311508 | -2.44734883 |
| EPHX3 | 4.105547081 | 2.20501E-06 | 4.035939236 | 2.65407E-05 | 4.439321229 | 8.2286E-06 | 4.193602515 |
| EVC2 | 2.149911458 | 0.016044474 | 2.446789036 | 0.006421237 | 2.073536041 | 0.022021542 | 2.223412178 |
| FADS2 | -1.266345312 | 0.001980602 | -1.0450584 | 0.021397385 | -1.038549642 | 0.018084179 | -1.11665112 |
| FAM133A | 3.575479503 | 0.000245832 | 2.980938054 | 0.005679079 | 3.320474245 | 0.001126305 | 3.292297268 |
| FAM78A | -2.507556489 | 7.37949E-07 | -1.807355461 | 0.001186411 | -2.223506738 | 1.88619E-05 | -2.1794729 |
| FAT3 | -2.486936086 | 1.70098E-08 | -1.295439388 | 0.01609397 | -1.262846598 | 0.016114565 | -1.68174069 |
| FBP1 | 2.900038012 | 4.76772E-06 | 2.555574121 | 0.000215055 | 2.763173468 | 4.17376E-05 | 2.7395952 |
| FBXL13 | 1.566478355 | 0.005113837 | 1.691975351 | 0.004070983 | 1.729929997 | 0.002459925 | 1.662794568 |
| FBXW10 | 3.528161726 | 1.98654E-09 | 3.531828396 | 1.54896E-08 | 3.416412442 | 7.81215E-08 | 3.492134188 |
| FLNC | 2.011680469 | 6.36545E-17 | 2.412913301 | 1.84596E-24 | 2.056044407 | 1.5592E-17 | 2.160212726 |
| FNDC4 | 1.501337854 | 0.002528902 | 1.575631657 | 0.002280339 | 1.423243296 | 0.0052321 | 1.500070936 |
| FUT3 | 1.642340467 | 8.62259E-06 | 1.004139582 | 0.026064693 | 1.319736028 | 0.001132936 | 1.322072026 |
| FYB1 | 2.900876466 | 3.25327E-05 | 2.291659398 | 0.004058903 | 2.223792577 | 0.00401978 | 2.47210948 |
| FZD4 | -1.015884519 | 0.002604579 | -1.415537846 | 9.0515E-06 | -1.134893132 | 0.000570145 | -1.18877183 |
| GCNA | 4.125633171 | 4.69919E-08 | 3.301837972 | 6.47901E-05 | 3.421047158 | 2.7173E-05 | 3.616172767 |
| GGT1 | 2.714702381 | 6.99816E-08 | 1.781194081 | 0.002357995 | 1.905644734 | 0.000625934 | 2.133847065 |
| GLIPR1 | 1.821967762 | 4.54416E-07 | 1.38316471 | 0.00052208 | 1.496209591 | 9.30052E-05 | 1.567114021 |
| GOLGA2P8 | -1.404180197 | 0.042050179 | -1.818546818 | 0.003808527 | -1.323121868 | 0.042789515 | -1.51528296 |
| GOLT1A | 1.285882762 | 0.00028933 | 1.016651385 | 0.012789718 | 1.025168144 | 0.008850054 | 1.109234097 |
| GPR173 | -1.019416979 | 0.026470862 | -1.04393766 | 0.027782652 | -1.126610192 | 0.011381174 | -1.06332161 |
| GSDMC | 1.570853557 | 0.000641371 | 1.736627461 | 0.000181335 | 1.595357149 | 0.000698738 | 1.634279389 |
| GSTA4 | -1.934122983 | 1.94072E-05 | -1.13894232 | 0.035293475 | -1.749428256 | 0.000181258 | -1.60749785 |
| GYG2 | -1.602162937 | 8.60728E-07 | -1.138263119 | 0.001256307 | -1.527312859 | 2.85376E-06 | -1.42257964 |
| H19 | -6.075822621 | 2.85984E-06 | -2.651242295 | 0.032270623 | -3.900468859 | 0.000195988 | -4.20917792 |
| HDAC9 | 1.649008377 | 1.21646E-07 | 1.334566822 | 8.04878E-05 | 1.503191839 | 3.37289E-06 | 1.495589013 |
| IL11 | 2.136472268 | 5.20904E-06 | 2.755134079 | 3.72149E-09 | 2.174393222 | 7.88486E-06 | 2.35533319 |
| IL31RA | 1.829149652 | 0.000208183 | 1.284816283 | 0.017996181 | 1.794169084 | 0.000188643 | 1.636045007 |
| IL6 | 1.969829411 | 1.81199E-05 | 1.506779737 | 0.003468091 | 1.5282522 | 0.002471328 | 1.668287116 |
| ITGA2 | 2.066020313 | 3.76651E-08 | 1.452258028 | 0.000532736 | 1.826753266 | 2.83673E-06 | 1.781677202 |
| KCNK6 | 1.010107157 | 0.005801723 | 1.297742716 | 0.000406933 | 1.298954585 | 0.000355289 | 1.202268153 |
| KILH | 1.678603043 | 9.41188E-15 | 1.684416519 | 1.23765E-13 | 1.495155982 | 9.21436E-11 | 1.619391848 |
| KPNA7 | 1.533805864 | 0.001202506 | 1.286172326 | 0.019388287 | 1.518266045 | 0.003741167 | 1.446081412 |
| KRT13 | 1.467128596 | 0.010290893 | 1.80912738 | 0.001329744 | 2.625223112 | 2.35156E-07 | 1.967159696 |
| KRT17 | 1.832141951 | 0.004318535 | 1.724116832 | 0.012313727 | 1.78350317 | 0.00660994 | 1.779920651 |
| KRT5 | 2.938201957 | 9.85731E-06 | 2.808201715 | 4.2406E-05 | 3.018564962 | 1.29042E-05 | 2.921656211 |
| KRT6A | 4.591774348 | 2.03873E-07 | 4.979458681 | 3.83734E-08 | 5.093837071 | 5.75116E-08 | 4.8883567 |
| KYNU | 1.841818527 | 3.30937E-11 | 1.080115155 | 0.000783513 | 1.530073096 | 1.38729E-07 | 1.484002259 |
| LAMA4 | 2.096680226 | 3.34592E-07 | 1.682886721 | 0.000179987 | 2.197993825 | 1.34389E-07 | 1.992520257 |
| LAMC2 | 1.538523772 | 8.75301E-14 | 1.277506986 | 2.91808E-09 | 1.239986621 | 8.17027E-09 | 1.352005793 |
| LAT2 | 3.334182211 | 4.16182E-08 | 2.480223682 | 0.000265547 | 2.735040739 | 4.1162E-05 | 2.849815544 |
| LINC01605 | 2.50434956 | 0.006388066 | 2.480045081 | 0.0130429 | 2.097791505 | 0.038905328 | 2.360728715 |
| LINC01914 | -1.432421334 | 0.018079638 | -1.480758084 | 0.015279647 | -1.587787115 | 0.007089147 | -1.50032218 |
| LIPG | 1.782759903 | 2.77289E-05 | 1.195323925 | 0.017494705 | 1.638162797 | 0.000209257 | 1.538748875 |
| LNCAROD | 1.560638846 | 0.003026733 | 1.691816774 | 0.001562655 | 1.324482003 | 0.017834424 | 1.525645874 |
| LRP2 | -7.234998549 | 0.00189867 | -5.205186868 | 0.021174372 | -6.8268655 | 0.00061673 | -6.42235031 |
| LRP4 | -1.562959914 | 4.56117E-06 | -1.15270884 | 0.002568351 | -1.459876959 | 3.37948E-05 | -1.39184857 |
| LRRC2 | 2.175671763 | 0.001016657 | 2.292659257 | 0.000664845 | 2.122523575 | 0.00200138 | 2.196951532 |
| LYPD6B | -1.476418961 | 4.35137E-05 | -1.306691607 | 0.000726696 | -1.277060596 | 0.000737424 | -1.35339039 |
| MAG | -1.872093227 | 3.17879E-06 | -1.38003484 | 0.002886359 | -1.420765337 | 0.001336852 | -1.55763113 |
| MAP2K6 | -1.435533242 | 0.001326419 | -1.29170931 | 0.006429788 | -1.326305643 | 0.004014986 | -1.35118273 |
| MAPT | -1.837947929 | 0.000223011 | -1.312088646 | 0.011899353 | -1.968525674 | 1.05383E-05 | -1.70618742 |
| MBOAT1 | -1.318914544 | 1.3753E-05 | -1.248347691 | 8.73672E-05 | -1.654207689 | 1.34531E-08 | -1.40715664 |
| MFNG | 3.329164006 | 1.03161E-11 | 2.730440813 | 7.91496E-08 | 2.44447011 | 5.85027E-06 | 2.834691643 |
| MROH3P | 8.248327176 | 0.004023674 | 6.736934783 | 0.029239063 | 8.371312545 | 0.005162314 | 7.785524835 |
| MSS51 | 1.969405851 | 0.020374571 | 1.925002516 | 0.03849396 | 2.270922299 | 0.008884178 | 2.055110222 |
| MT1E | 6.291562718 | 1.21488E-06 | 5.105442027 | 0.000133965 | 4.422426144 | 0.000568692 | 5.27314363 |
| MUC4 | 2.973684592 | 7.28615E-05 | 2.834216234 | 0.000377183 | 3.372530949 | 8.07125E-06 | 3.060143925 |
| MYH10 | -2.054211582 | 1.3372E-06 | -1.61655908 | 0.000463324 | -2.358843191 | 1.13697E-08 | -2.00987128 |
| MYL7 | -4.722863762 | 3.98885E-05 | -3.184490145 | 0.012624461 | -4.547100418 | 5.29284E-05 | -4.15148477 |
| N4BP2L1 | -1.065949337 | 0.024287664 | -1.038346776 | 0.037347444 | -1.209390055 | 0.008187825 | -1.10456206 |
| NALCN | 1.37314595 | 0.01866889 | 1.290958781 | 0.044465314 | 1.707549895 | 0.002414294 | 1.457218209 |
| NAV3 | 1.167890352 | 6.50035E-07 | 1.634453502 | 7.15953E-13 | 1.607189968 | 1.67598E-12 | 1.469844607 |
| NCCRP1 | 1.906385612 | 0.000575811 | 1.893523698 | 0.00164954 | 2.220471666 | 0.000102787 | 2.006793659 |
| NECTIN4 | 1.549060122 | 0.026409282 | 1.759666815 | 0.013332651 | 1.767496761 | 0.009661486 | 1.692074566 |
| NEGR1 | 1.969135603 | 0.000389392 | 1.75744604 | 0.003750371 | 1.839149652 | 0.001366994 | 1.855243765 |
| NELL2 | -1.946767804 | 0.002901307 | -1.87828685 | 0.004636512 | -1.946835381 | 0.001600983 | -1.92396334 |
| NLGN1 | -1.889329304 | 0.027091877 | -2.525635337 | 0.001981671 | -2.613041665 | 0.000309258 | -2.34266877 |
| NLRP1 | 3.471823665 | 0.006046593 | 2.857313955 | 0.035844507 | 2.709476688 | 0.033915347 | 3.012871436 |
| NLRP4 | 5.718214637 | 0.006445404 | 5.573622302 | 0.012528098 | 6.411532159 | 0.002229045 | 5.901123033 |
| NOS3 | 4.829624837 | 3.99428E-08 | 4.23100986 | 1.95232E-05 | 3.246749872 | 0.00085303 | 4.102461523 |
| NOTUM | -2.5067664 | 4.55271E-05 | -2.056897927 | 0.001855952 | -2.95158657 | 6.4331E-07 | -2.50508363 |
| NPNT | -3.451541812 | 1.47917E-12 | -2.440502751 | 1.47683E-06 | -3.045429419 | 1.1623E-10 | -2.97915799 |
| NPY4R | -1.878341135 | 2.30672E-05 | -1.373280134 | 0.005679079 | -1.707129426 | 0.000187171 | -1.6529169 |
| OR7E14P | 2.239393239 | 0.041321089 | 2.431076854 | 0.043732408 | 2.708884888 | 0.02023319 | 2.459784994 |
| PCDH18 | -2.286769282 | 3.15971E-08 | -1.911213345 | 2.16206E-06 | -2.588890111 | 1.23331E-11 | -2.26229091 |
| PDE2A | 3.117745559 | 1.56699E-05 | 3.211932342 | 4.86596E-05 | 3.367361125 | 2.06771E-05 | 3.232346342 |
| PEAR1 | 1.264246044 | 0.002492314 | 1.284733011 | 0.004058903 | 1.663037169 | 6.15953E-05 | 1.404005408 |
| PHGDH | -1.135267632 | 1.47917E-12 | -1.007036552 | 8.5185E-10 | -1.418436042 | 8.55563E-20 | -1.18691341 |
| PKDCC | -1.481651644 | 4.18411E-09 | -1.071001447 | 9.12095E-05 | -1.109624104 | 3.30652E-05 | -1.22075906 |
| PLAC8 | -1.373933163 | 2.11776E-07 | -1.071058798 | 0.000232586 | -1.17682334 | 2.591E-05 | -1.20727177 |
| PLAUR | 1.492320491 | 4.39749E-05 | 1.448521695 | 0.000171987 | 1.352766149 | 0.00042754 | 1.431202778 |
| PLBD1 | -1.301791207 | 1.00325E-08 | -1.002502197 | 3.68187E-05 | -1.310559321 | 6.54624E-09 | -1.20495091 |
| PPP1R1C | 1.61904062 | 0.000277669 | 1.421963561 | 0.003699511 | 1.133189015 | 0.023959966 | 1.391397732 |
| PPP1R9A | 1.188279669 | 0.012430563 | 1.325505916 | 0.006204111 | 1.339812746 | 0.004310227 | 1.284532777 |
| PPP4R3C | 5.13381308 | 0.002870963 | 4.478781787 | 0.019005296 | 5.426022761 | 0.002436477 | 5.012872543 |
| PRICKLE1 | -1.749236897 | 1.41673E-05 | -1.259025553 | 0.005656886 | -1.582356394 | 0.000123509 | -1.53020628 |
| PRSS21 | 1.352980186 | 0.00061503 | 1.782073512 | 2.22167E-06 | 1.366962483 | 0.000737424 | 1.50067206 |
| PTK6 | 1.319543607 | 0.000302243 | 1.059879683 | 0.012528098 | 1.273039876 | 0.001346058 | 1.217487722 |
| PTPRB | 1.74955921 | 1.61991E-05 | 1.366589382 | 0.003153926 | 1.32367114 | 0.003059101 | 1.479939911 |
| PTPRH | 2.047975843 | 1.00325E-08 | 1.646023651 | 4.86596E-05 | 1.626206885 | 8.38144E-05 | 1.773402126 |
| PURPL | 2.610865781 | 0.001134132 | 2.061295083 | 0.021379112 | 3.032723852 | 0.000144033 | 2.568294905 |
| PUS10 | 1.075335043 | 0.02076093 | 1.124122934 | 0.022059099 | 1.100358917 | 0.021270848 | 1.099938965 |
| RBMS3 | -1.068635785 | 0.040407497 | -1.284640483 | 0.012789718 | -1.258210363 | 0.01187584 | -1.20382888 |
| RIPOR2 | 3.896104531 | 0.002688888 | 3.466882821 | 0.021908876 | 3.780515916 | 0.014247187 | 3.714501089 |
| RND1 | 2.170596516 | 0.001313615 | 1.790315176 | 0.02658363 | 2.023365854 | 0.00956654 | 1.994759182 |
| RPS6KL1 | 2.263949296 | 0.000111066 | 2.505208421 | 2.92578E-05 | 2.052872116 | 0.001101522 | 2.274009944 |
| SAMD4A | 1.225863798 | 0.001166135 | 1.393005243 | 0.000233137 | 1.183890571 | 0.002229045 | 1.267586537 |
| SBK3 | 2.050080524 | 0.010340731 | 2.452488088 | 0.002954915 | 2.557078429 | 0.0013941 | 2.35321568 |
| SEC14L2 | 2.056286391 | 1.17408E-25 | 1.579383526 | 1.23765E-13 | 1.845076071 | 2.71179E-18 | 1.826915329 |
| SELPLG | 2.256675697 | 0.024115759 | 2.545492876 | 0.015369433 | 2.249504736 | 0.030385976 | 2.35055777 |
| SERPINB7 | 1.876890503 | 4.58071E-05 | 1.514533018 | 0.003404587 | 1.763591333 | 0.000263886 | 1.718338285 |
| SERPINE1 | 1.752487218 | 2.68669E-26 | 1.257327556 | 5.48271E-13 | 1.346469697 | 3.04962E-15 | 1.452094824 |
| SLC13A3 | 2.784468106 | 1.0384E-05 | 2.401902108 | 0.000540395 | 2.07025142 | 0.004186349 | 2.418873878 |
| SLC22A13 | 3.253571297 | 0.010780816 | 3.006912392 | 0.039738278 | 3.488999461 | 0.013637958 | 3.249827717 |
| SLC2A3 | -1.86364871 | 0.006734724 | -2.367166307 | 0.000384388 | -2.159171281 | 0.001224425 | -2.12999543 |
| SLC30A4-AS1 | 2.079180723 | 0.015356076 | 2.252666225 | 0.015023173 | 2.193598955 | 0.01801452 | 2.175148634 |
| SLC37A2 | 2.485611531 | 3.04096E-06 | 1.930789199 | 0.001861256 | 2.894441369 | 2.35156E-07 | 2.436947366 |
| SLC44A4 | 4.154825423 | 2.13102E-10 | 3.345499285 | 1.41779E-05 | 5.406278078 | 3.53007E-13 | 4.302200929 |
| SLC7A7 | -1.973342885 | 1.19297E-06 | -1.282362142 | 0.006410734 | -1.347551537 | 0.002746554 | -1.53441885 |
| SLC7A8 | -2.41027423 | 0.000344436 | -1.831364052 | 0.01288651 | -2.748802275 | 2.51265E-05 | -2.33014685 |
| SMAD6 | -1.485638071 | 1.11229E-06 | -1.360102257 | 1.95232E-05 | -1.768588661 | 3.10627E-09 | -1.53810966 |
| SMPDL3A | -1.509441166 | 0.013027614 | -1.342083657 | 0.044465314 | -1.351026123 | 0.032575778 | -1.40085032 |
| SOCS1 | -1.810944744 | 0.002586275 | -2.066174602 | 0.000407653 | -2.137194626 | 0.000181258 | -2.00477132 |
| SOX9 | 1.01536728 | 0.041389679 | 1.892656778 | 1.8465E-05 | 1.416646013 | 0.002287677 | 1.44155669 |
| SPOCK1 | 2.022203162 | 0.012512352 | 2.408355409 | 0.00267787 | 1.95415766 | 0.017509739 | 2.128238744 |
| SPTBN5 | 1.992609859 | 2.82149E-06 | 2.51241041 | 1.78843E-09 | 1.810795391 | 0.000118566 | 2.105271887 |
| STEAP1 | 1.857432815 | 0.008127152 | 2.134862931 | 0.004123877 | 1.992896667 | 0.010142547 | 1.995064138 |
| STX1A | 1.582529339 | 6.54552E-09 | 1.780600544 | 1.27359E-10 | 1.323235755 | 7.31716E-06 | 1.562121879 |
| SUSD2 | -1.514139222 | 0.003685755 | -1.492367157 | 0.006229058 | -2.129954664 | 1.2644E-05 | -1.71215368 |
| SYNPO | 1.96753634 | 0.000839741 | 1.404751294 | 0.038998832 | 1.646879421 | 0.006896855 | 1.673055685 |
| SYT16 | 1.742727654 | 0.000432483 | 1.705681618 | 0.001268143 | 1.610934532 | 0.002327104 | 1.686447935 |
| TAC3 | -3.770906665 | 1.37503E-09 | -2.359562583 | 0.000523762 | -3.087556278 | 7.9741E-07 | -3.07267518 |
| TCIM | 1.4650116 | 0.001631414 | 1.087763859 | 0.043957864 | 1.046531396 | 0.044039787 | 1.199768951 |
| TET1 | -1.616082122 | 2.32529E-05 | -1.114486738 | 0.009986554 | -1.264381376 | 0.001896175 | -1.33165008 |
| TGFA | 1.375534968 | 1.4299E-08 | 1.182333803 | 4.2126E-06 | 1.403025048 | 9.49744E-09 | 1.32029794 |
| TGFBR3 | -1.743354904 | 0.000626849 | -1.311653249 | 0.020902998 | -1.652937935 | 0.001336852 | -1.56931536 |
| TLE4 | -4.453596758 | 5.65914E-08 | -3.506838072 | 1.05041E-05 | -3.514204491 | 7.88486E-06 | -3.82487977 |
| TMEM140 | 2.254103763 | 0.001806506 | 2.251051091 | 0.003468091 | 2.002655912 | 0.008879131 | 2.169270255 |
| TMEM171 | 1.575481245 | 0.006712756 | 1.609408864 | 0.007897145 | 1.763654673 | 0.002339196 | 1.649514927 |
| TMEM37 | -2.165094803 | 0.007274105 | -2.088435255 | 0.014552485 | -1.956664329 | 0.022021542 | -2.0700648 |
| TMEM52B | -1.117557865 | 0.002472042 | -1.30365469 | 0.00038218 | -1.419447493 | 5.46998E-05 | -1.28022002 |
| TMPRSS2 | -2.384276038 | 7.4794E-05 | -1.645360573 | 0.017782105 | -1.978306512 | 0.001914784 | -2.00264771 |
| TMT1A | -1.702193219 | 1.51829E-06 | -1.677459685 | 3.70666E-06 | -1.653164374 | 4.59575E-06 | -1.67760576 |
| TNFAIP3 | 1.662057254 | 5.43335E-05 | 1.720099598 | 5.72157E-05 | 1.579424848 | 0.000226976 | 1.653860567 |
| TRBC2 | 1.780021691 | 0.004201952 | 2.619730745 | 6.92035E-06 | 2.300994379 | 0.000101155 | 2.233582272 |
| TRPV1 | 1.18426186 | 0.003599314 | 1.09460856 | 0.01403436 | 1.045897774 | 0.016724734 | 1.108256065 |
| TTC28 | -1.038135795 | 0.031441632 | -1.068693431 | 0.028944116 | -1.302901196 | 0.003505305 | -1.13657681 |
| VAV1 | 1.781539266 | 3.36712E-11 | 1.913291557 | 6.75809E-12 | 1.810396978 | 1.23489E-10 | 1.835075934 |
| VIM | 1.735107042 | 0.043170314 | 2.644839605 | 0.00077979 | 2.046379473 | 0.012788182 | 2.142108706 |
| VIT | -5.853643969 | 0.000523275 | -4.362528376 | 0.00574197 | -6.152851905 | 1.88253E-05 | -5.45634142 |
| VNN1 | 2.381600011 | 5.21898E-05 | 2.255502965 | 0.000894146 | 3.424054151 | 3.03685E-08 | 2.687052375 |
| VSIG2 | 2.107022449 | 0.011642303 | 2.245526529 | 0.011990475 | 2.645925878 | 0.001651798 | 2.332824952 |
| WNK4 | 1.380448445 | 0.004075878 | 1.136834539 | 0.045420289 | 1.375834819 | 0.007377414 | 1.297705934 |
| XYLT1 | 7.042600857 | 0.000161481 | 6.616727598 | 0.001191129 | 6.138902308 | 0.002851322 | 6.599410254 |
